# Characterising group-level brain connectivity: a framework using Bayesian exponential random graph models

**DOI:** 10.1101/665398

**Authors:** B.C.L. Lehmann, R.N. Henson, L. Geerligs, Cam-CAN, S.R. White

## Abstract

The brain can be modelled as a network with nodes and edges derived from a range of imaging modalities: the nodes correspond to spatially distinct regions and the edges to the interactions between them. Whole-brain connectivity studies typically seek to determine how network properties change with a given categorical phenotype such as age-group, disease condition or mental state. To do so reliably, it is necessary to determine the features of the connectivity structure that are common across a group of brain scans. Given the complex interdependencies inherent in network data, this is not a straightforward task. Some studies construct a group-representative network (GRN), ignoring individual differences, while other studies analyse networks for each individual independently, ignoring information that is shared across individuals. We propose a Bayesian framework based on exponential random graph models (ERGM) extended to multiple networks to characterise the distribution of a entire population of networks. Using resting-state fMRI data from the Cam-CAN project, a study on healthy ageing, we demonstrate how our method can be used to characterise and compare the brain’s functional connectivity structure across a group of young individuals and a group of old individuals.

## 1 Introduction

Brain connectivity analysis aims to understand how spatially distinct regions of the brain interact with each other. The use of networks to model whole-brain connectivity has become increasingly popular in recent years [5]: by treating distinct regions as nodes and the connections between them as edges, researchers have gained new insights into both the structure and function of the brain. To determine the salient features of the brain’s connectivity structure, it is necessary to identify which of those are common across individuals. Given the complex interdependencies inherent in network data, however, it is not a trivial task to consider connectivity structure across multiple individuals; how best to combine connectivity information across participants remains a key challenge.

Many existing methods aim to construct a (single) group-representative network (GRN) across (multiple) individuals. There have been several methods proposed for constructing GRNs. For example, Achard et al. [2] constructed a group-representative network using mean functional connectivity (mean-GRN) by first taking the mean of the individuals’ functional connectivity matrices and then thresholding the resulting matrix. Song et al. [38] took a similar approach using the median of the individuals’ functional connectivity matrices (i.e. median-GRN). Sinke et al. [36] constructed a group-representative structural connectivity network from Diffusion Tensor Imaging (DTI) data by keeping those edges which are present in at least 35% of the individuals’ networks (i.e. minimal-GRN). These edge-based GRN methods are computationally convenient, since each individual is processed separately and the subsequent analysis is based on a single network. However, these methods ignore higher-order topological properties present in each individual’s network. Rather that construct a completely new network as a combination of each individual’s, other methods represent a whole group by selecting a ‘best’ individual network in terms of mutual information [25] or Jaccard index [20] (i.e. best-GRN). While this approach preserves the topological properties of a single network, in subsequent analyses it may lead to undue weight being placed on features that are not present in many of the remaining networks.

The above issues are compounded when comparing brain networks across groups of individuals, identifying group differences in connectivity via differences in the GRNs from each group. In particular, network metrics are influenced by the overall density of the network [17]. As a result, it is difficult to disentangle differences in more complex topological properties such as clustering from differences in the network density; especially in the context of the various GRN methods. This is particularly important in the context of ageing since mean functional connectivity is known to decrease with age [15]. Note that it is not trivial to simply control for network density when comparing network metrics due to their highly non-linear relationship.

A key consideration is whether to view the network as a single object or a single realisation of a random process. The former implies that the network is a fixed construct, the latter defines a random process (with inherent noise) that gives rise to a distribution of brain networks. An exponential random graph model (ERGM) defines a parametric statistical distribution across all possible networks with a given set of nodes (see [30] for a review). The aim of the model is to characterise the distribution of a network in terms of a set of *summary statistics*. These summary statistics are typically comprised of topological features of the network, such as the number of edges and subgraph counts. The summary statistics enter the likelihood via a weighted sum; the weights are (unknown) model parameters that quantify the relative influence of the corresponding summary statistic on the overall network structure and must be inferred from the data. ERGMs are thus a flexible way in which to describe the global network structure as a function of network summary statistics.

ERGMs have been applied successfully to characterise both functional connectivity [34, 33, 27, 11] and structural connectivity [36]. To date, there have been two proposed approaches for using ERGMs in group studies. The first approach constructs an edge-based GRN for each group and then fits the same ERGM (i.e. identical summary statistics) separately to each group’s network, obtaining GRN-ERGM parameter estimates [36]. As described above, this edge-based approach may average over informative topological structure present in each of the individuals’ networks. In contrast, the second approach fits an ERGM to each individuals’ network and then takes the mean or median of the fitted parameters from individuals within a group to represent the group-level connectivity structure [33, 27] (i.e. mean-ERGM or median-ERGM). Note that it is then possible to generate a GRN from the resulting mean-ERGM and median-ERGM parameter estimates [33]. While this is preferable to the first approach, taking a simple mean or median may obscure important information.

In this article, we introduce a novel framework to model a population of individual brain networks which we call multi-BERGM (the ‘B’ denotes Bayesian; the framework could theoretically be analysed in a frequentist paradigm but we do not consider that further). Our framework uses a Bayesian formulation of the ERGM within a multi-level (i.e hierarchical) model, which allows us to pool information across multiple individuals nested within multiple groups. This provides an approach to characterise connectivity at a group-level, and thus compare the brain’s connectivity structure between groups. We demonstrate our method on functional connectivity networks derived from resting-state fMRI scans of participants from the Cam-CAN project, a study on healthy ageing [32].

## 2 Methods

### 2.1 Data

The data were collected as part of Phase II of the CamCAN project (www.cam-can.org; [32]). The MRI data were acquired on a 3T Siemens TIM Trio at the MRC Cognition & Brain Sciences Unit, with a 32 channel head-coil. Structural images were acquired using a 1mm3 isotropic, T1-weighted Magnetization Prepared RApid Gradient Echo (MPRAGE) sequence and a 1mm3 isotropic, T2-weighted Sampling Perfection with Application optimized Contrasts using different flip angle Evolution (SPACE) sequence. The fMRI data for the eyes-closed, resting-state run were acquired using a Gradient-Echo (GE) Echo-Planar Imaging (EPI) sequence, and consisted of 261 volumes (lasting 8 min and 40 s). Each volume contained 32 axial slices (acquired in descending order), with slice thickness of 3.7 mm and interslice gap of 20% (for whole brain coverage including cerebellum; TR 1,970 ms; TE 30ms; voxel-size 3 mm x 3 mm x 4.44 mm). EPI fieldmaps with two TEs (5.19ms and 7.65ms) were also acquired. All the raw data from CamCAN Phase II, together with more acquisition details, are available on: http://www.cam-can.org/index.php?content=dataset.

The data were processed using r7219 of the SPM12 software (http://www.fil.ion.ucl.ac.uk/spm), automated with release r5.4 of the Automatic Analysis (AA) pipeline system [10] (http://github.com/automaticanalysis/automaticanalysis/releases/v5.4.0; for overview of pipelines, see [40]) in r2015 of MATLAB (The MathWorks). To obtain a good starting-point for image normalisation, the T1 image was coregistered to the Montreal Neurological Institute (MNI) template using rigid body transformation, and then the T2 image was coregistered to the T1. Both T1 and T2 images were bias corrected, and then combined in a multimodal segmentation to estimate images of each of six tissue classes, including grey-matter (GM), white-matter (WM) and cerebrospinal fluid (CSF). Diffeomorphic registration (DARTEL) was then applied to the GM and WM segments to create group templates, which were in turn transformed to MNI space using a 12-parameter affine transform.

The fMRI images were unwarped using distortion fields estimated from the fieldmaps, and corrected for motion using rigid-body realignment to the mean fMRI image across runs. The different slice acquisition times were corrected by interpolating to the middle slice. The images were rigid-body coregistered to the T1 image and the spatial transformations from that T1 to MNI space (diffeomorphic and affine) were applied to every fMRI image. Residual effects of abrupt motion were reduced by applying wavelet despiking [28]. The mean time series for all voxels within the thresholded WM and CSF segments were calculated to use as later covariates of no interest.

The fMRI time series were then extracted from 90 cortical and subcortical regions of interest (ROIs) from the AAL atlas [42]. ROIs in the cerebellum were not included. The time series for each voxel in each ROI were adjusted for various confounds by taking the residuals from a general linear model (GLM) that included: 1) the time series in WM and CSF segments, 2) the 6 rigid-body motion parameters from the realignment stage, 3) the first-order difference in those motion parameters across successive TRs and 4) a cosine basis set up to cut-off frequency of 0.008Hz (implementing a form of high-pass filter). Second-order (squared) terms for 1), 2) and 3) were also included in the GLM. The autocorrelation in the GLM error was modelled by a family of 8 exponentials with half-lives from 0.5 to 64 TRs (estimated by pooling across voxels within each ROI), and used to prewhiten the time series. The average voxels of the residual time series was then used as a summary measure for each ROI. This approach was based on the optimised pipeline proposed by Geerligs et al. [15].

To illustrate our method, we analyse two groups (denoted *j*, i.e. *j* = 1, 2) from the Cam-CAN study: the 100 youngest individuals, aged 18-33, and the 100 oldest individuals, aged 74-89 (denoted *i* within each group, i.e. *i* = 1, …, *n*_*j*_, where *n*_1_ = *n*_2_ = 100).

### 2.2 Network construction

The preprocessed data consists of, *N* = 90, ROI time series for each individual. To apply a standard (binary) ERGM, it is necessary to extract a network for each individual. For a given individual *i*, we first computed the pairwise Pearson correlation between each of the time series, yielding a *N* × *N* correlation matrix ***C***^(*i*)^. In contrast with the pipeline proposed by Geerligs et al. [15], we did not apply mean regression to the resulting functional connectivity matrices because we accounted for differences in mean connectivity via the thresholding procedure or through the specification of the ERGM (we return to this point in the discussion).

To each correlation matrix, we then applied a threshold *r*^(*i*)^ to produce an *N* ×*N* adjacency matrix, ***A***^(*i*)^, with entries:

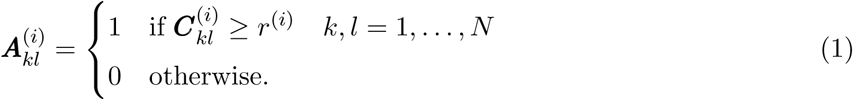

The adjacency matrix defines an individual’s network, ***y***^(*i*)^, with an edge between nodes *k* and *l* if and only if 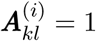. The choice of threshold *r*^(*i*)^ can have a significant impact on the resulting connectivity structure [43, 18]. The two most common strategies are “absolute” thresholding and “proportional” thresholding. Absolute thresholding picks a common threshold *r*^(*i*)^ = *r* for each individual. In contrast, proportional thresholding ensures that the same number of edges are present in each network by allowing the threshold to vary by individual.

Both thresholding strategies have their drawbacks. Since the number of edges in a network inherently affects the overall network [43, 16], differences in other network metrics may simply be attributable to variations in the overall connectivity. A proportional threshold may thus be preferred in order to keep the number of edges constant across individuals (in both groups) and facilitate comparison of other metrics of interest. On the other hand, a lower correlation value may be less reliable in indicating a functional connection between brain regions. Therefore, by including lower correlations as edges for individuals with lower overall connectivity, a proportional threshold may induce more randomness in the resulting network [18].

Given these issues, we analysed networks constructed using the two different thresholding procedures: absolute and proportional. Further, for both procedures we considered two distinct threshold values (i.e. different *r*^(*i*)^s), chosen to yield average node degrees of *K* = 3 and *K* = 5. The value of *K* = 3 corresponds to the efficiency-cost optimization (ECO) criterion [13]; this aims to optimise the trade-off between the efficiency of a network and its density, or wiring cost. The value of *K* = 5 has been used in previous applications of ERGMs to resting-state functional connectivity [34, 33].

#### 2.2.1 Group-representative network construction

In order to contrast our framework with existing methods, we also constructed group-representative networks (GRN) following Achard et al. [2] (mean-GRN) and Song et al. [38] (median-GRN). The mean-GRN and median-GRN take the mean and median, respectively, of the correlation matrices across all individuals within the group (young and old) and then threshold these group mean and median matrices to yield networks with average node degrees of *K* = 3 and *K* = 5. Explicitly, 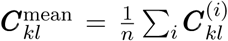 and 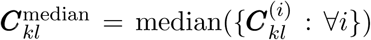, with associated adjacency matrices. These approaches produce a single network for each group, which can be modelled as an ERGM and we can compare the respective GRN-ERGM parameters.

### 2.3 Exponential random graph model specification

The probability mass function of a network ***Y*** under an ERGM is given by *π*(***y***|*θ*) where

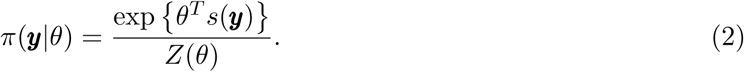

Here, *s*(***y***) is a vector of *p* network summary statistics, *θ* ∈ Θ ⊆ ℝ^*p*^ is a vector of *p* corresponding model parameters that must be estimated from the data and *Z*(*θ*) =Σ_***y*′**∈*𝒴*_exp {*θ*^*T*^ *s*(***y*′**)} is the normalising constant ensuring the probability mass function sums to one. Given data, that is, an observation ***y*** of ***Y***, the goal is to infer which values of *θ* best correspond to the data under this distribution. The summary statistics included in a given ERGM represent those configurations expected to appear more frequently or less frequently than in a random graph. In other words, the choice of summary statistics is a modelling decision; it reflects our belief of how the global network structure may be summarised and is driven by the context of the network. The flexibility of ERGMs derives from the range and number of possible summary statistics that can be used [29].

In what follows, we work in the Bayesian paradigm, treating the model parameters *θ* as random variables. The Bayesian formulation of ERGMs was first suggested by Koskinen [21] and then expanded upon by Caimo & Friel [7]. Through the machinery of Bayesian hierarchical modelling, this provides a natural framework for describing the joint distribution of a *group* of networks. To fully specify a Bayesian ERGM, we need only augment the definition in (2) with a prior distribution *π*(*θ*) for the model parameters. Given an observation ***y*** of the network, we can then perform inference by analysing the posterior distribution *π*(*θ*|***y***). Through Bayes’ rule, we may write the posterior as

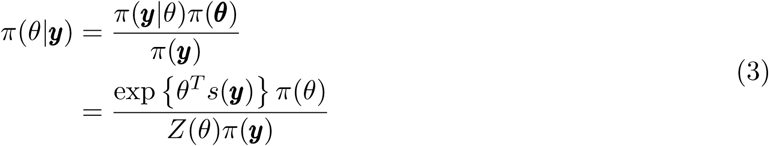

where *π*(***y***) = ∫_Θ_ *π*(***y***|*θ*)*π*(*θ*)*dθ* is the model evidence. Although the posterior distribution is generally not available in a closed-form expression, one may obtain samples from the posterior through a Markov chain Monte Carlo method known as the exchange algorithm [26].

For the absolute thresholding our exponential random graph model (which is common to all individuals and groups) uses three summary statistics; these have previously been applied to single brain functional connectivity networks [34, 35]. The three summary statistics used are: number of edges (E); geometrically weighted edgewise shared partners (GWESP); and geometrically weighted non-edgewise shared partners (GWNSP). For the proportional thresholding, we fixed the number of edges and modelled the network using two summary statistics: GWESP and GWNSP.

The number of edges *E* = Σ_*k*<*l*_ *Y*_*kl*_ characterises the sparsity of the network. When using an absolute threshold, it is important to include this term in the model in order to account for differences in the network density across individuals. When using a proportional threshold, however, we opted not to include this summary statistic in the model, as the number of edges is fixed across individuals. In this case, we constrained the space of networks to those with the same number of edges as those observed. In other words, the model describes a probability distribution over networks with a given average node degree *K*. Note that this in contrast to previous applications of ERGMs in neuroimaging, which include the number of edges in the model despite using a proportional threshold [34, 33].

The geometrically weighted edgewise shared partner (GWESP) statistic of a network ***y*** is a measure of network transitivity (or clustering) and is given by:

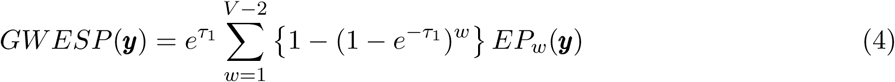

where *EP*_*w*_(***y***) is the number of *connected* node pairs having exactly *w* shared partners and *τ*_1_ > 0 is a decay parameter. The decay parameter *τ*_1_ serves to diminish the effect of the network having more higher-order edgewise shared partners relative to lower-order edgewise shared partners. In other words, the increase in GWESP from adding a single edge is smaller if the edge adds a shared partner to connected nodes that already share many partners.

The geometrically weighted non-edgewise shared partner (GWNSP) statistic is similarly defined as

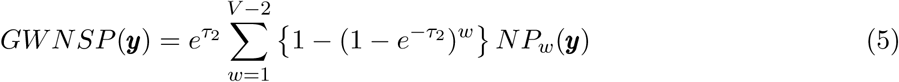

where *NP*_*w*_(***y***) is the number of *non-connected* node pairs having exactly *w* shared partners and *τ*_2_ > 0 is a decay parameter. GWNSP is related to global network efficiency; a higher value of GWNSP indicates that non-connected nodes are more likely to have a shared partner.

While it is possible to treat the decay parameters for both geometrically weighted statistics as extra model parameters (leading to *curved* ERGMs [19]), this increases the computational burden substantially. The decay parameters were therefore fixed at *τ*_1_ = *τ*_2_ = 0.75, as these values generally resulted in better fitting models [34].

### 2.4 Multi-level (hierarchical) framework

The Bayesian exponential random graph model described above provides a flexible family of distributions for a *single* network. Our goal is to extend this to a model for a *population* of networks. Our proposed approach is simple: represent each network as a separate ERGM within a Bayesian multilevel (or hierarchical) model. By pooling information across individual networks, this approach allows us to characterise the distribution of the whole population.

We model each individual network ***Y*** ^(*i*)^ as an exponential random graph with model parameter *θ*^(*i*)^. Importantly, each individual ERGM must consist of the same set of summary statistics *s*(·). The probability mass function of each network can then be written

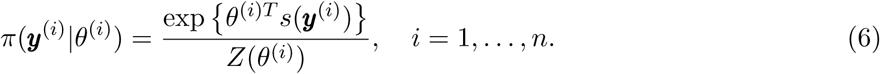

This specifies the data-generating process for each individual network. To obtain a joint distribution for the set of networks, we assume that, conditional on their respective individual-level parameters, the ***Y*** ^(*i*)^ are independent. Thus, the sampling distribution for the set of networks ***Y*** is simply the product of the individual probability mass functions:

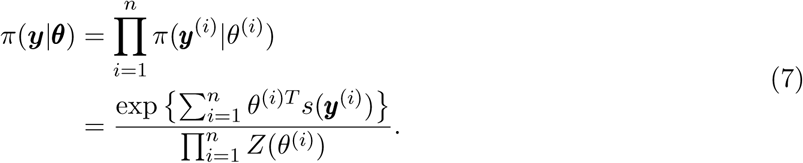

#### First level of the multi-level model (within group)

To model a group of networks we need to specify the prior distribution of the individual-level ERGM parameters 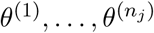 (for individuals 1, …, *n*_*j*_ in group *j*). To this end, we propose a multilevel model such that *θ*^(1)^, …, *θ*^(*n*)^ are drawn from a common population-level Normal distribution with parameters *ϕ* = (*µ*_multi_, **Σ**_*θ*_), which are also treated as random variables. We write

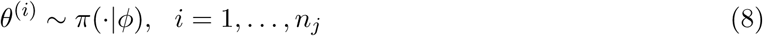

for the population-level distribution. Assuming that, conditional on *ϕ*, the *θ*^(*i*)^ are independent, we have

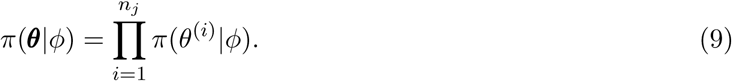

Finally, write *π*(*ϕ*) for the (hyper)prior distribution of *ϕ*. The joint distribution of (***Y***, ***θ***, *ϕ*) can be written as *π*(***y, θ***, *ϕ*) = *π*(***y***|***θ***)*π*(***θ***|*ϕ*)*π*(*ϕ*). See Figure 1 for a diagrammatic representation of the full hierarchical framework. Full details of the prior specifications can be found in the Supplementary Material.

**Figure 1:**
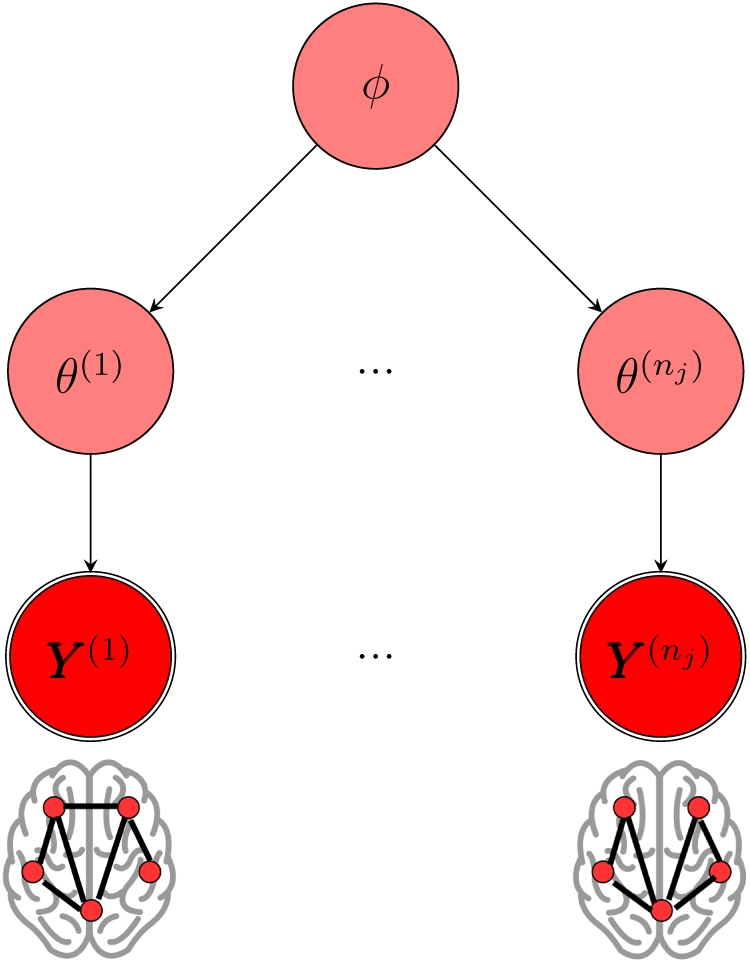
A diagrammatic representation of the hierarchical framework. Each network ***Y*** ^(*i*)^ is modelled as an exponential random graph with individual-level parameter *θ*^(*i*)^. In turn, each *θ*_*i*_ is assumed to come from a common population-level distribution with hyperparameter *ϕ*.

#### Second level of the multi-level model (between group)

To extend the model for multiple groups of within a population, we add another level to the hierarchy. Denoting *θ*^(*i,j*)^ to be the (vector-valued) ERGM parameter for the *i*^*th*^ individual in the *j*^*th*^ group, we assume 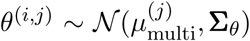. Thus, we assume different group-level means but the same covariance structure across groups (our framework allows per-group covariance, but the sample size of our illustrative example leads us to a common covariance). Writing 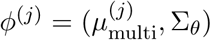, we then complete the model by specifying the group-level prior *ϕ*^(*j*)^ ∼ *π*(·|*ϕ*^*pop*^) and population-level hyperprior *π*(*ϕ*^*pop*^). Full details of the prior specifications can be found in the Supplementary Material.

### 2.5 Model fitting and comparison to alternative approaches

To generate samples from the joint posterior distribution, we devised a novel MCMC algorithm, the details of which can be found in (Lehmann, 2019) [24]. We used our algorithm to generate posterior samples for two sets of networks. The first set consisted of a single group of networks from the 100 youngest individuals in the Cam-CAN study (a one-level model), while the second set also included the networks from the 100 oldest individuals (a two-level model).

#### One-level model for young group

For the single-group dataset, we derived estimates for the posterior mean and 95% credible regions for each pair of components of the group-level parameter *µ*_multi_ using our multi-BERGM model with one-level (i.e. individuals nested within one group).

In order to compare our method with existing approaches, we also generated posterior samples for the same (using three summary statistics defined earlier) Bayesian ERGM fit to the mean-GRN, *µ*_mean-GRN_, and median-GRN, *µ*_median-GRN_.

We also compare our method, which generates a full posterior distribution for *µ* and Σ, to a mean-BERGM approach. Specifically, we fit a BERGM to each individual within the group (i.e. we fit *n*_1_ = 100 separate single-BERGMs and obtain 100 separate sets of posterior samples 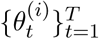 for *i* = 1, …, 100). The original proposal of this approach used frequentist ERGMs, giving a point estimate per individual. The Bayesian approach we use yields a separate posterior per individual, which can then be combined into a group-level estimate 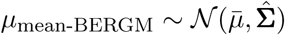 with mean

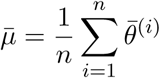

and covariance

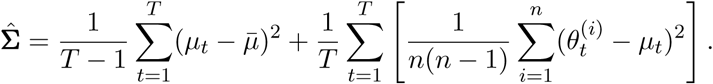

where 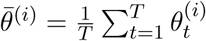 and 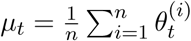.

#### Two-level model for young and old groups

For the two-group dataset, we used the two-level multi-BERGM model and derived posterior density estimates for the group-level mean parameters 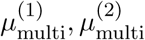, thus providing a way to compare the connectivity structure across the two groups. We also applied the mean-BERGM approach to the group of old individuals, i.e. we fit a single-BERGM to each individual within the old group and used the resulting posterior samples to construct a group-level estimate (see previous section).

### 2.6 Goodness-of-fit assessment

For the single-group dataset, we assessed the goodness-of-fit of the posterior distribution to the data by simulating networks from the posterior predictive distribution and comparing the network metrics of these simulated networks to those of the observed networks. Specifically, we chose uniformly at random *S* = 1000 samples *µ*_1_, …, *µ*_*S*_ generated from the posterior distribution of the group-level parameter *µ*_multi_. For each sample, *s*, we also randomly selected (uniformly, with replacement) an individual *i*_*s*_ and then simulated a network ***Y*** ^(*s*)^ from the ERGM with parameter 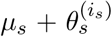. We compared the simulated networks to the observed networks based on three network metrics: degree distribution, geodesic distance distribution (length of shortest paths) and edge-wise shared partners distribution (note these are different to, but highly correlated with, the summary statistics used to define the ERGM).

It is possible to assess goodness-of-fit on any network metrics by comparing simulated networks to observed networks. Since age-related differences in local efficiency [1, 14, 37, 31] and global efficiency [1, 31] have previously been observed, we assessed the goodness-of-fit for the two-group dataset in terms of these two metrics. For both groups, we chose uniformly at random *S* = 1000 samples 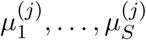 generated from the posterior distribution of the group-level parameter 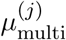. For each sample, we simulated a network ***Y*** ^(*j,s*)^ from the ERGM with parameter 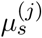, and compared the local and global efficiency of the simulated networks to the observed networks.

### 2.7 Availability of data and code

Access to the Cam-CAN dataset or the purpose of scientific investigation, teaching or the planning of clinical research studies can be requested at https://camcan-archive.mrc-cbu.cam.ac.uk/dataaccess/datarequest.php. The code to fit the Bayesian multilevel framework with exponential random graph models is available to download as an R package at https://github.com/brieuclehmann/multibergm. The scripts used to generate the results for this paper are available at https://doi.org/10.6084/m9.figshare.11919072.v1.

## 3 Results

We performed analyses for two populations of networks, the first with a single group (young only) and the second with two groups (young and old). Both the one and two group settings were considered under two different thresholding procedures (absolute and proportional) and two different threshold values (chosen to yield average node degrees of *K* = 3 and *K* = 5); resulting in eight sets of networks that we analysed using our multi-BERGM framework. In this section we present the results of the analyses performed on the sets of networks constructed under absolute thresholding with a population-wide average node degree of *K* = 3. The other results can be found in the Supplementary Material.

In the context of this paper the supplementary results are in line with those presented here, except where explicitly stated.

For the single-group setting we compare our method, specifically the estimated ERGM parameters, to several alternative approaches from the literature.

### Single-group mean ERGM parameter estimation

We first consider the point estimate of the ERGM parameters, *µ*, for the young only (i.e. single-group setting) under various approaches: our one-level multi-BERGM framework; the mean-BERGM, the mean-GRN BERGM; the median-GRN BERGM; and the distribution of point estimates (posterior sample mean) from a single-BERGM fitted to each individual’s network. Briefly, multi-BERGM fits all the model parameters simultaneously in a multilevel framework; mean-BERGM combines estimates from the separate single-BERGM fits; mean-GRN and median-GRN are single-BERGM fits on the group-representative mean and median networks respectively (see Section 2.5 for details).

Figure 2 presents the three components of *µ* (corresponding to the three summary statistics) showing the mean-BERGM estimate (blue dotted line); the posterior sample means of the one-level multi-BERGM (red line), the mean-GRN BERGM (green dashed line), and the median-GRN BERGM (purple dashed line); and the distribution of posterior sample means for single-BERGM fitted to each individual (grey histogram).

**Figure 2:**
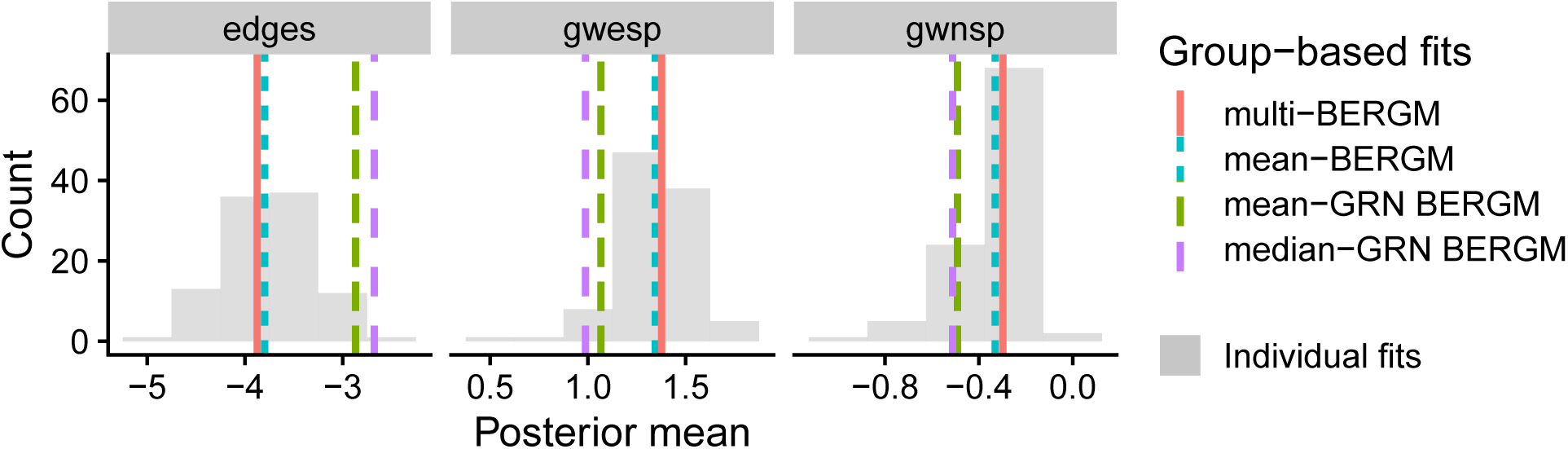
Posterior mean estimates of models fit to networks constructed under absolute thresholding with group-wide average node degree *K* = 3. The histogram (grey bars) corresponds to the single-BERGM posterior mean values of the model parameters fitted to each of the *n* = 100 individual networks. The vertical lines correspond to the posterior mean values of group-level mean parameter under multi-BERGM (red line), mean-BERGM (blue dotted line), mean-GRN BERGM (green dashed line) and median-GRN BERGM (purple dashed line). For each of the model components, the posterior mean under multi-BERGM lay close to the mean-BERGM estimate, i.e. the centre of the single-BERGM mean values. In contrast, for the edges and GWESP components, posterior means for the mean-GRN BERGM and median-GRN BERGM lay in the tails of the single-BERGM mean values.

The multi-BERGM estimates are very close to the mean-BERGM estimates, i.e. the centre of the posterior mean values across the individual network fits (single-BERGMs). In contrast, the posterior means of the edges and GWESP components for the mean-GRN and median-GRN are in the tails of single-BERGM distribution. Similar results were observed under the proportional thresholding procedures while the estimates under absolute thresholding with average node degree *K* = 5 were similar under each of the approaches (see Supplementary Material Figure A.1). This suggests that the mean-GRN and median-GRN do not always accurately capture the topological structure of the majority of the individual networks.

### Individual-level covariance under multi-BERGM and single-BERGM

We consider the covariance of the posterior distribution of the individual-level parameters for three randomly selected participants under our multi-BERGM approach compared to the distribution from the single-BERGM fits. For each approach, we calculated 95% credible regions for each pair of model components based on the respective posterior samples.

While the mean values were similar between the multi-BERGM and single-BERGM distributions, the multi-BERGM yielded tighter credible regions (Figure 3). This illustrates one of the benefits of modelling the networks simultaneously: the borrowing of information across individuals leads to more precise estimates of the model parameters. The credible regions derived from the other thresholding procedures were similar or tighter under multi-BERGM compared to single-BERGM (see Supplementary Material Figure A.2).

**Figure 3:**
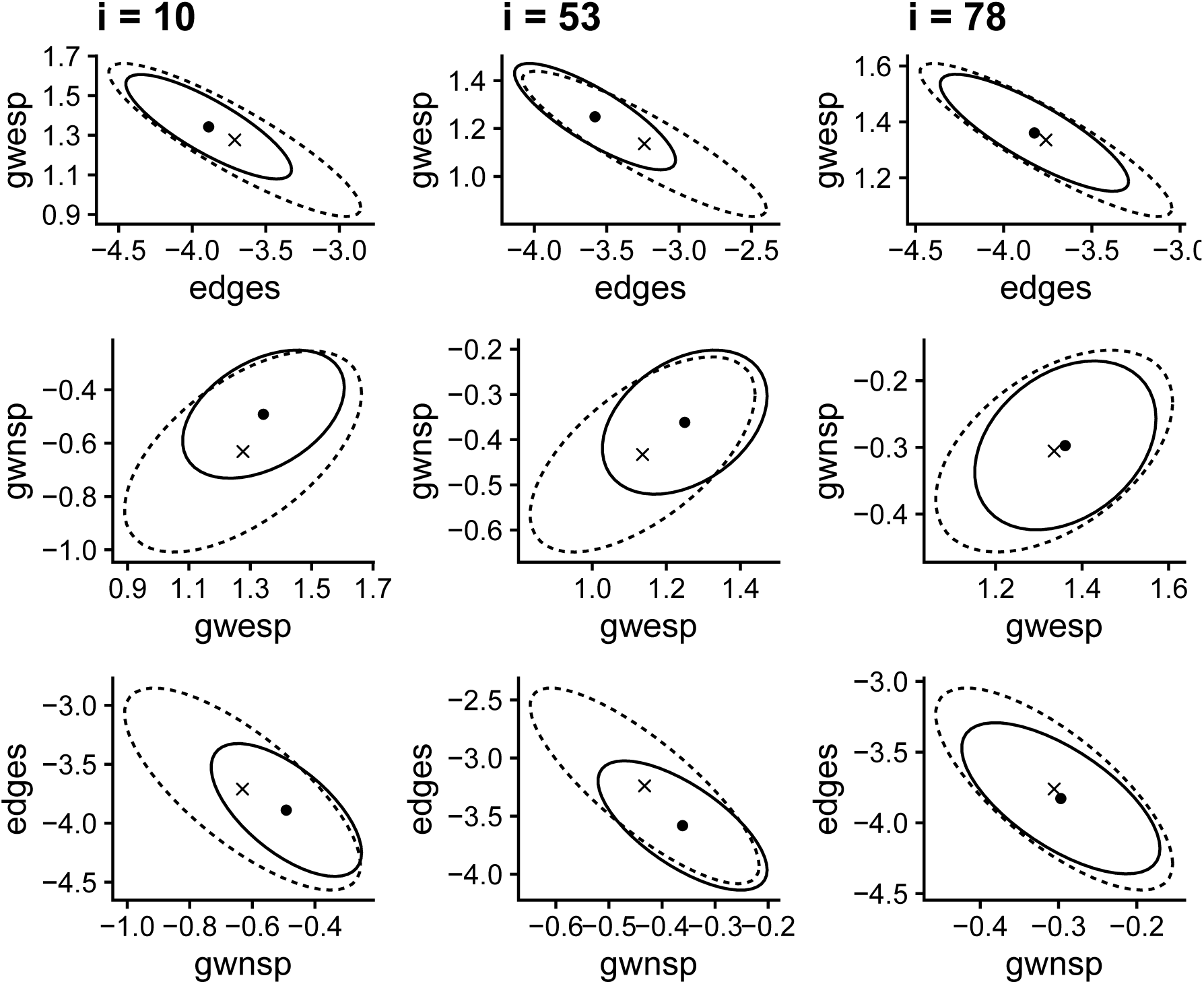
Posterior means and 95% credible regions of the individual-level parameters for three subjects and for each pair of model components under the multi-BERGM (dot and solid line) and under the single-BERGM (cross and dashed line) approaches. The correlation between parameter estimates is shown by the elliptical shape of the regions.

### Single-group covariance under multi-BERGM and mean-BERGM

We consider the covariance of the posterior distribution under our multi-BERGM approach compared to the mean-BERGM distribution constructed from the posterior samples the single-BERGMs for each individual. From the posterior samples for the multi-BERGM, we calculated 95% credible regions for each pair of model components. For the mean-BERGM approach, we calculate a 95% region from a normal distribution with mean and covariance based on the single-BERGM fits for each individual.

While the mean values were similar between the multi-BERGM and mean-BERGM distributions, the multi-BERGM yielded moderately tighter credible regions (Figure 4). This again illustrates how the borrowing of information across individuals can lead to more precise estimates of the model parameters. The credible regions derived from the other thresholding procedures were similar or tighter under multi-BERGM compared to the mean-BERGM approach (see Supplementary Material Figure A.3).

**Figure 4:**
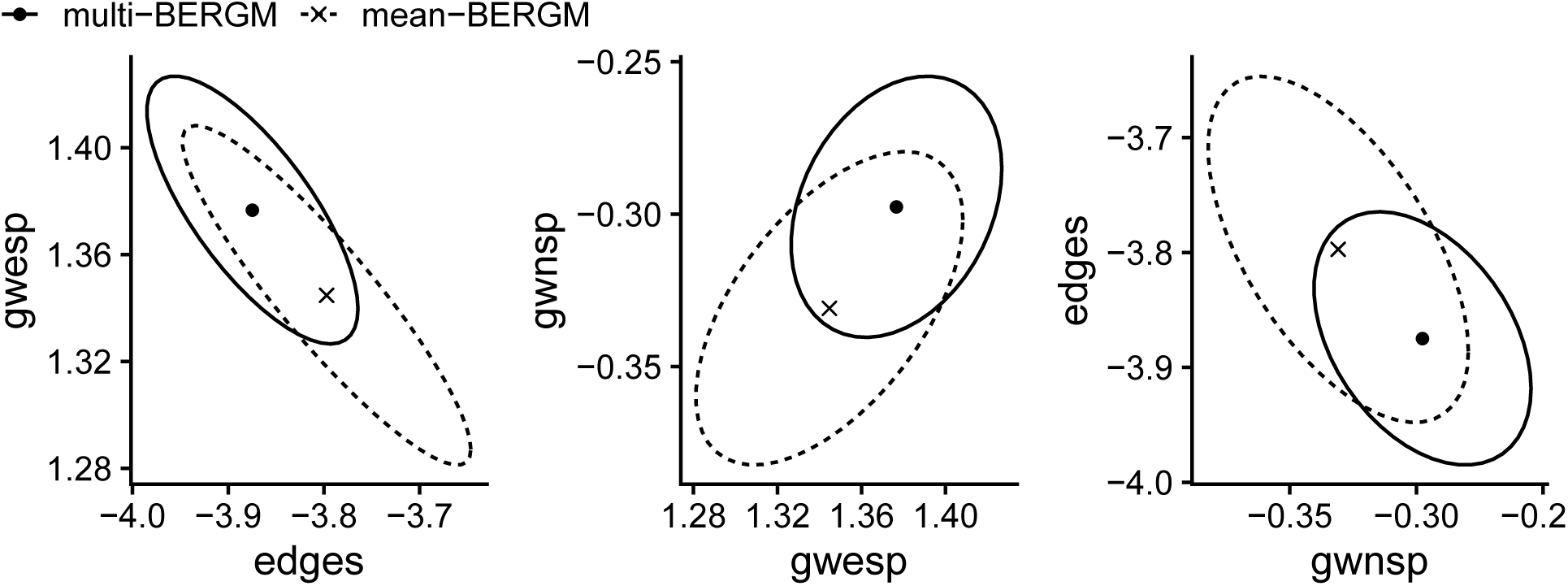
Posterior means and 95% credible regions of the group-level mean parameters for each pair of model components under the multi-BERGM (dot and solid line) and under the mean-BERGM (cross and dashed line) approaches.

### Single-group goodness-of-fit under multi-BERGM

To assess the goodness-of-fit under multi-BERGM, we simulated networks from the ERGM with parameters taken uniformly at random from the posterior samples. The simulated networks appeared broadly similar to the observed networks in terms of the network metrics we considered: degree distribution, geodesic distance distribution and edgewise-shared partner distribution (Figure 5). This indicates that the group-level posterior distribution adequately captured the important network characteristics across the entire group of individuals. Similar results were observed under the remaining three thresholding procedures (see Supplementary Material Figures A.4-A.6).

**Figure 5:**
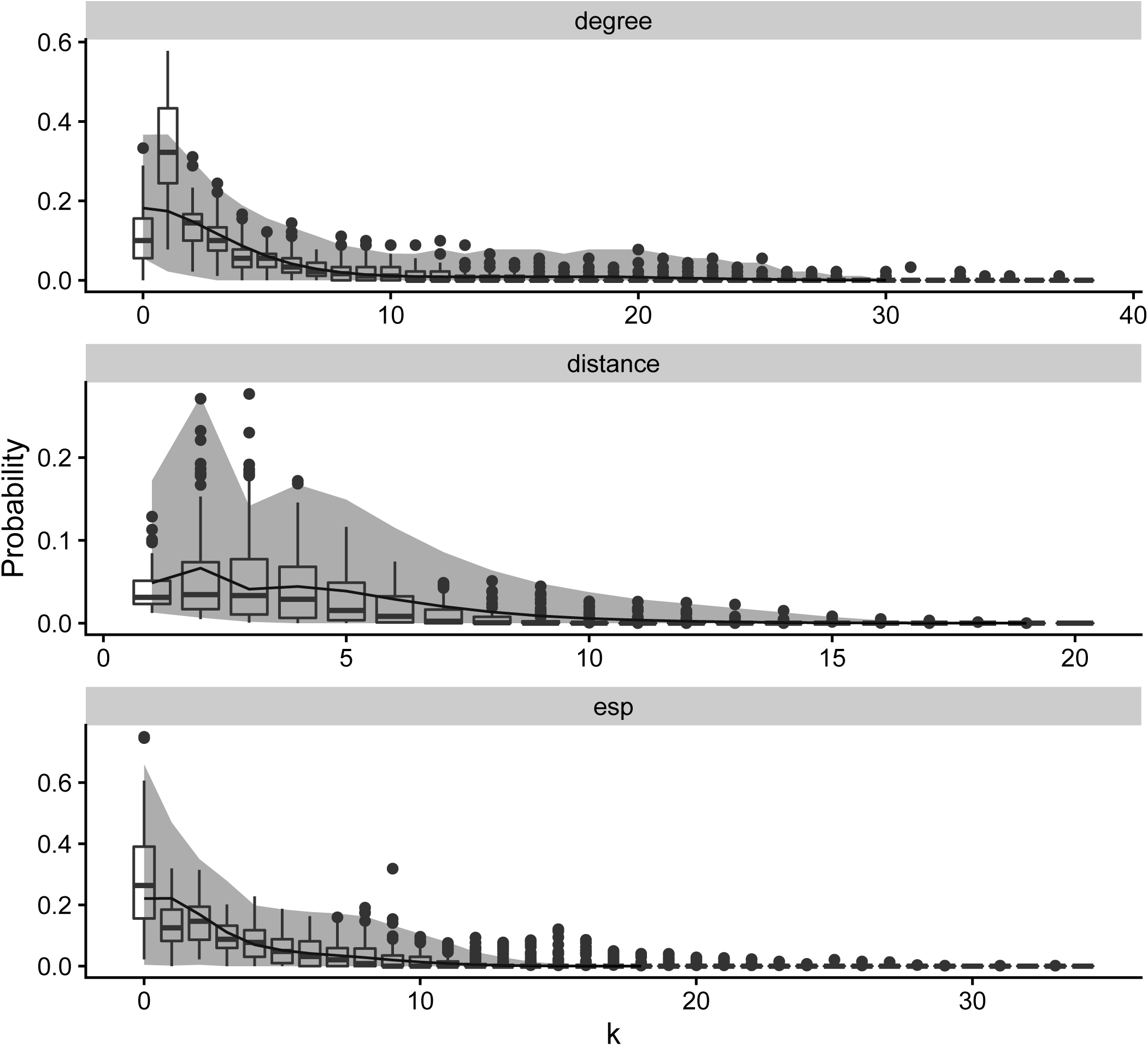
Goodness-of-fit assessment of the multi-BERGM for a single group. *S* = 1000 networks were generated from the ERGM with model parameters sampled from the respective group-level distributions. The simulated networks are compared to the observed data (shown in the box plots) across three network metrics: degree distribution, geodesic distance distribution, edge-wise shared partner distribution. The lines correspond to the respective means across the simulated networks, while the ribbons correspond to 95% credible intervals.

### Two-group multi-BERGM: group differences

Using a two-level multi-BERGM framework we obtain posteriors for the group-level ERGM parameters. These posterior distributions reveal that network differences between the young group and the old group are driven by differences in the GWNSP parameter (Figure 6). In particular, the posteriors for the edges and GWESP parameters are almost identical for the two groups, while the GWNSP parameter is markedly larger for the young group. This difference in the group-level GWNSP parameters was also observed under the remaining three thresholding procedures (Figures A.7-A.8).

**Figure 6:**
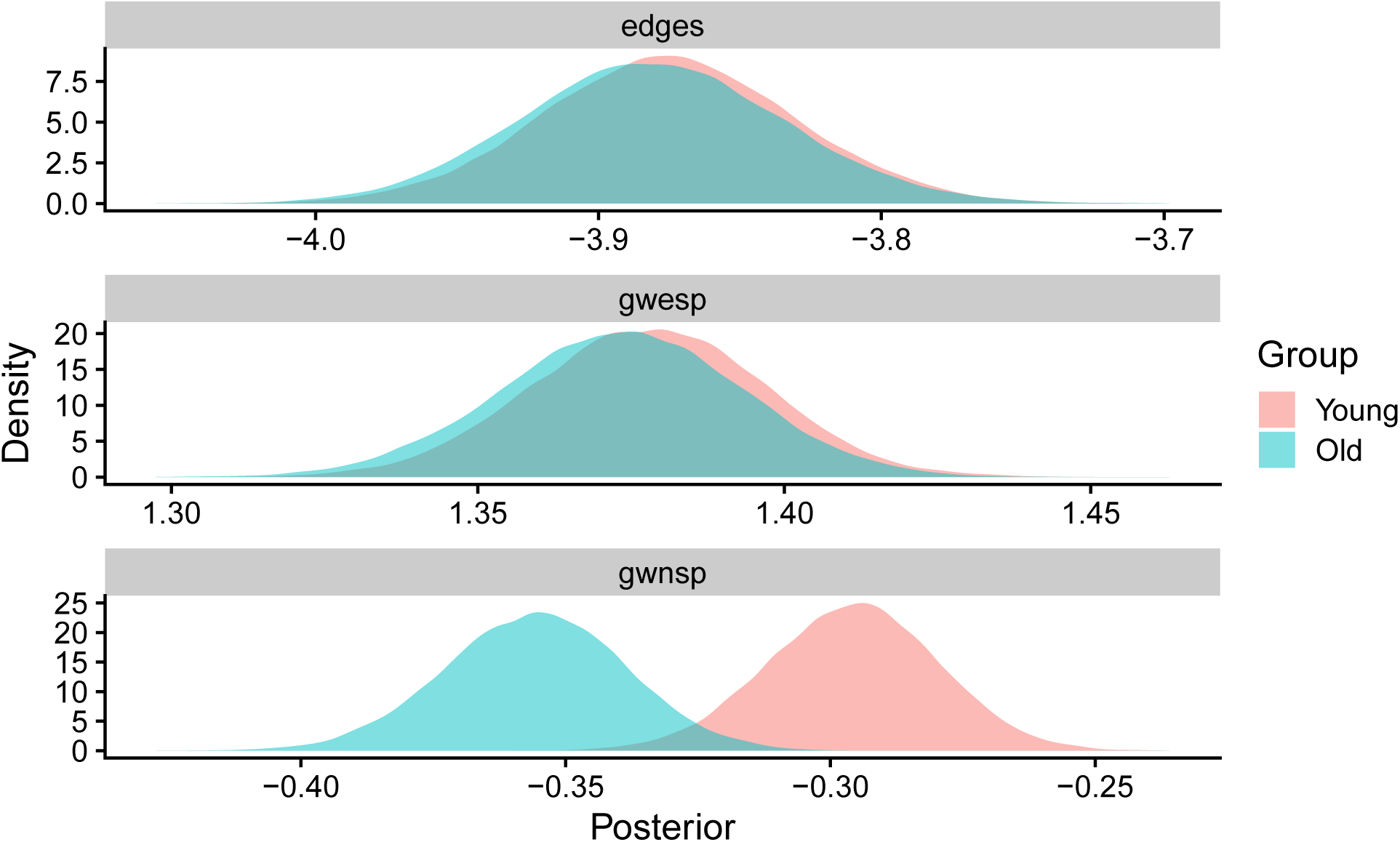
Posterior density estimates under multi-BERGM for the group-level mean parameters for networks constructed via absolute thresholding with average node degree *K* = 3. The young group displayed markedly larger values for the GWNSP parameter while the edges and GWESP posteriors were almost identical between the two groups.

By considering the posterior distribution of difference in the group-level parameters, *µ*_1_ − *µ*_2_, we also calculated 95% credible regions to assess the degree of certainty in the group differences under both the multi-BERGM and the mean-BERGM approaches (Figure 7). The left panel shows that difference in the (edges-GWESP) pair of parameters is centered on zero, while the middle and right panels demonstrate that, under the multi-BERGM approach, the credible regions for the (GWESP-GWNSP) and (GWNSP-edges) pair do not contain zero. This provides evidence that the group-level differences in network structure are driven by the differences in GWNSP.

**Figure 7:**
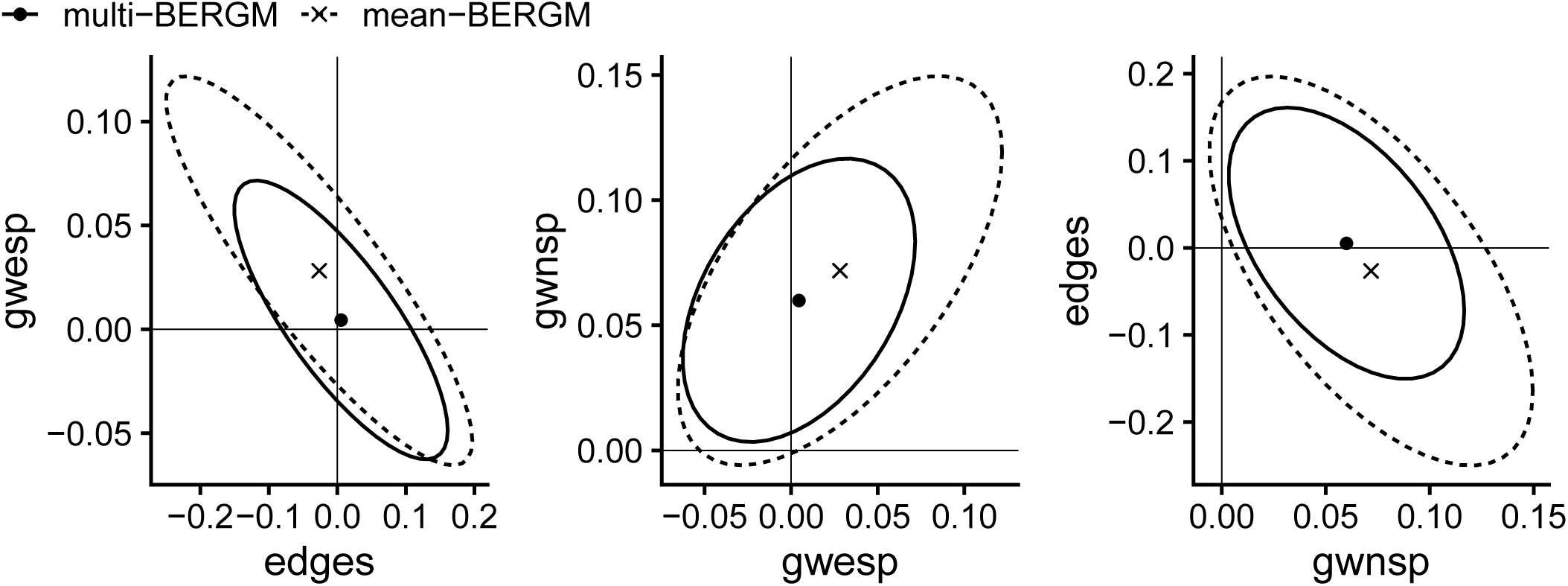
Posterior means and 95% credible regions of the difference in group-level mean parameters for each pair of model components under the multi-BERGM (dot and solid line) and under the mean-BERGM (cross and dashed line) approaches.

Under the mean-BERGM approach, the credible regions are slightly wider and contain zero for each pair of parameters. This again highlights the benefit of more precise estimates of the model parameters via the multi-BERGM approach by modelling the networks simultaneously. Similar results were observed under absolute thresholding with average node degree *K* = 5 but no significant differences were found under proportional thresholding (Figure A.10)

Age-related differences in both local efficiency [1, 14, 37, 31] and global efficiency [1, 31] have previously been observed. To check whether the multi-BERGM approach could identify similar age-related group differences, we also simulated *S* = 1000 networks for each group from the ERGM with parameters taken uniformly at random from the group-level posterior samples. Overall, the simulated networks for the young group exhibited higher global and local efficiency, in correspondence with the observed data (Figure 8). Similar results were observed for networks constructed via absolute thresholding with average node degree *K* = 5 (Supplementary Figure A.11). Under proportional thresholding, however, the posterior predictive distribution of local and global efficiency did not correspond well with the observed networks, indicating a poor model fit (Supplementary Figures A.12-A.13).

**Figure 8:**
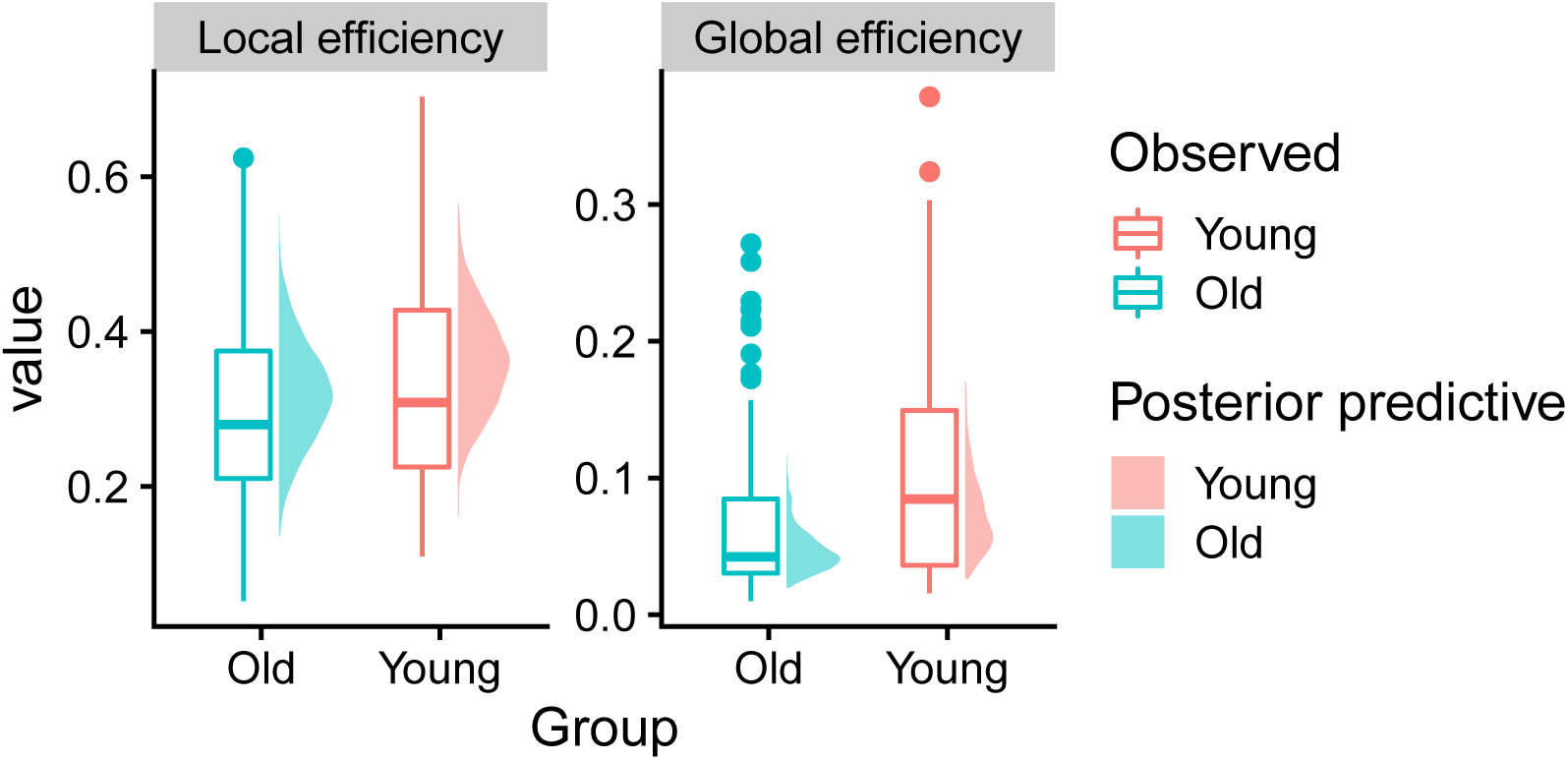
Local and global efficiency in the observed networks (bar plots) compared to *S* = 1000 networks simulated from the posterior predictive distribution (density plots) for the young group (red) and the old group (blue).

Based on the results from previous sections, we have shown that the various GRN methods do not properly reflect the distribution of individual ERGMs, and hence we do not consider group differences under these alternative approaches.

## 4 Discussion

The characterisation of connectivity structure at a group level is a necessary step towards understanding how the brain changes with a given phenotype such as age, disease condition or mental state. We have proposed multi-BERGM, a multi-level (hierarchical) Bayesian framework based on exponential random graph models (ERGMs), as a coherent method for understanding group level connectivity. By pooling information across the individual networks, this framework provides a principled (and extendable) approach to model the relational structure for groups of networks.

Our work builds on an previous approaches attempting to investigate differences between groups of networks, either by constructing a group-representative network (GRN) [2, 38, 36, 25, 20] or by combining separate individual-level estimates, e.g. mean-ERGM [33, 27]. There are three main advantages of our framework: using the Bayesian paradigm allows us to formally quantify uncertainty in the group-level model parameters; the group-level posterior densities provide a way to directly compare connectivity structure across groups; and the multi-level (hierarchical) framework allows information to be borrowed across individuals thus, in principle, improving the accuracy of the parameter estimates.

Our proposed method takes as input any group of networks. While there are a variety of methods to construct networks from functional neuroimaging data, each with their own benefits and disadvantages, we took the most common and simple approach of applying a threshold to the correlation matrices, keeping edges between nodes that exhibited a sufficiently strong positive correlation. The choice of threshold can have a significant impact on the resulting connectivity structure [43, 18] and we found that different threshold values did indeed lead to small differences in the posterior distribution of the model parameter. Regardless of the threshold used, our approach provides a framework to characterise the group-level brain connectivity structure, though further investigation into the effect of thresholding procedure on ERGM parameter estimates is warranted. Our framework could theoretically be used with weighted ERGMs [12, 44], thus bypassing the thresholding issue, though the associated computational cost is likely to be prohibitive.

From a modelling perspective, the appropriate selection of summary statistics is crucial. This is true of fitting ERGMs to single networks, let alone populations of networks. Moreover, the ‘correct’ choice likely depends on the network construction method. We used the summary statistics suggested in Simpson et al. [34], which were selected via a graphical goodness-of-fit approach, in which the ‘best’ metrics are chosen from a prespecified set of potential metrics. While it is possible to perform Bayesian model selection for an ERGM on a single network [6, 3], further work is needed to extend these approaches to perform model selection for a group of networks.

A graphical assessment of the model indicated a reasonable goodness-of-fit under constant thresholding. In particular, the model was able to recover differences in observed global and local efficiency, which were both higher in the young group than the old group. The observed age-related decrease in local efficiency is in line with previous studies [1, 14, 37, 31]. A decline in global efficiency has also been observed previously [1, 31], though not in other studies [14, 37]. In our study, the thresholding method had a pronounced effect on the observed age-differences. Under constant thresholding, we observed a clear difference in global efficiency between the two age groups and a relatively small difference in local efficiency. Under proportional thresholding, however, both local and global efficiency were slightly lower for the young group. The disparities between thresholding procedures may be associated with alterations in mean connectivity due to age-related differences in vascular health [15], while the discrepancies with previous studies are likely due to differences in network construction including the parcellation and definition of functional connectivity.

A downside to our framework, especially compared to the single-(B)ERGM and GRN approaches, is in terms of computation cost. Using a 40-core computing cluster (Intel Xeon E7-8860 v3, 2.2 GHz), the algorithm took between 8 and 12 hours to produce the posterior samples for each multi-BERGM (with additional time to tune the Bayesian procedure). However, Krivitsky & Handcock [23] provide empirical evidence suggesting the computational cost may grow on the order of *p*(*N* +*E*) log(*E*) where *p* is the number of summary statistics, *N* is the number of nodes, and *E* is the (typical) number of edges (as well as increase roughly linearly in the number of networks, *n*). In practical terms this limits the size of networks that our framework can currently handle to relatively coarse parcellations of the brain (approximately 100 regions). Improving the efficiency of fitting (B)ERGMs is an ongoing area of research [4, 39, 9] and our framework will benefit from ongoing developments and may be applicable to more spatially resolved networks in the future. The code used to apply our framework is available as an R package at https://github.com/brieuclehmann/multibergm.

The flexibility of Bayesian hierarchical modelling could be exploited to deal with more complex group structures such as factorial designs. Similarly, given multiple networks per individual, it would be (conceptually if not computationally) straightforward to extend the framework by adding another layer to the model. It would then be possible to apply our framework to dynamic functional connectivity or task-based scans in order to query how connectivity structure changes with time or across states.

## 4.1 Acknowledgements

BCLL is supported by the UK Engineering and Physical Sciences Research Council [Programme number EP/R018561/1] and Jesus College, Oxford. SRW and RNH are supported by the UK Medical Research Council [Programme numbers U105292687 and MC-A060-5PR10]. The Cambridge Centre for Ageing and Neuroscience (Cam-CAN) was supported by the Biotechnology and Biological Sciences Research Council (grant number BB/H008217/1). LG is funded by a Rubicon grant [446-13-013] and a Veni grant [451-16-013] from the Netherlands Organization for Scientific Research.

The Cam-CAN corporate author consists of the project principal personnel: Lorraine K Tyler, Carol Brayne, Edward T Bullmore, Andrew C Calder, Rhodri Cusack, Tim Dalgleish, John Duncan, Richard N Henson, Fiona E Matthews, William D Marslen-Wilson, James B Rowe, Meredith A Shafto; Research Associates: Karen Campbell, Teresa Cheung, Simon Davis, Linda Geerligs, Rogier Kievit, Anna McCarrey, Abdur Mustafa, Darren Price, David Samu, Jason R Taylor, Matthias Treder, Kamen Tsvetanov, Janna van Belle, Nitin Williams; Research Assistants: Lauren Bates, Tina Emery, Sharon Erzinlioglu, Andrew Gadie, Sofia Gerbase, Stanimira Georgieva, Claire Hanley, Beth Parkin, David Troy; Affiliated Personnel: Tibor Auer, Marta Correia, Lu Gao, Emma Green, Rafael Henriques; Research Interviewers: Jodie Allen, Gillian Amery, Liana Amunts, Anne Barcroft, Amanda Castle, Cheryl Dias, Jonathan Dowrick, Melissa Fair, Hayley Fisher, Anna Goulding, Adarsh Grewal, Geoff Hale, Andrew Hilton, Frances Johnson, Patricia Johnston, Thea Kavanagh-Williamson, Magdalena Kwasniewska, Alison McMinn, Kim Norman, Jessica Penrose, Fiona Roby, Diane Rowland, John Sargeant, Maggie Squire, Beth Stevens, Aldabra Stoddart, Cheryl Stone, Tracy Thompson, Ozlem Yazlik; and administrative staff: Dan Barnes, Marie Dixon, Jaya Hillman, Joanne Mitchell, Laura Villis.

## A Supplementary Material

### A.1 Prior specification

The choice of prior distributions for Bayesian ERGMs has yet to be studied in any great detail. The appropriate setting of priors is a challenging task due to the typically high levels of dependence between parameters [22]. Studies thus far have generally assumed (flat) multivariate normal prior distribution on the model parameters [7, 36, 41]. For simplicity, we also assume multivariate normal priors, though alternative specifications fully warrant further investigation.

#### Single group

For the single-group model, we assume

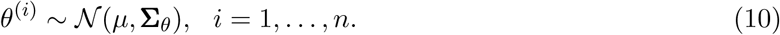

For the hyperprior distribution of *ϕ* = (*µ*, **Σ**_*θ*_), we assume a partially conjugate prior. This assumes *µ* and **Σ**_*θ*_ are a priori independent, placing a multivariate normal prior on *µ* and an inverse-Wishart prior on **Ψ**:

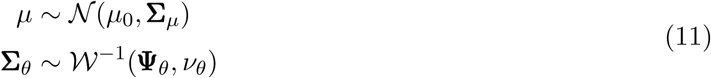

where *µ*_0_ ∈ ℝ^*p*^, *ν*_*θ*_ ∈ ℝ is such that *ν*_*θ*_ > *p*−1, and **Σ**_*µ*_, **Ψ**_*θ*_ are constant, *p* × *p* positive-definite matrices. These choices of hyperpriors are motivated by simplicity as well as computational convenience.

We set *ν*_*θ*_ = *p* and use an empirical approach to set the hyperparameters (*µ*_0_, **Σ**_*µ*_, **Ψ**_*θ*_). We do this by first fitting a single Bayesian ERGM to each network, using the `Bergm` package [8], with the same summary statistics as in the full multilevel model. For each network ***y***^(*i*)^, we produced *K* = 15000 samples 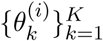 from their respective posterior distributions *π*(*θ*^(*i*)^|***y***^(*i*)^). In particular, we obtained estimates 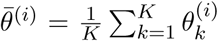 of the mean and 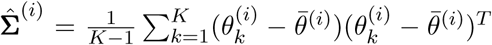 of the covariance for the posterior distribution from each network. Writing 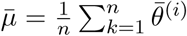, we then set 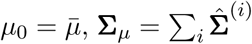 and 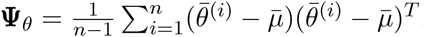.

#### Multiple groups

For the multiple-group model, we assume

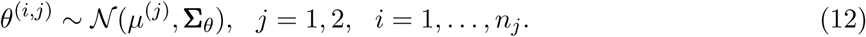

For the group-level prior distribution of *ϕ*^(*j*)^ = (*µ*^(*j*)^, **Σ**_*θ*_), we assume a partially conjugate prior. This assumes *µ*^(*j*)^ and **Σ**_*θ*_ are a priori independent, placing a multivariate normal prior on *µ*^(*j*)^ and an inverse-Wishart prior on **Ψ**:

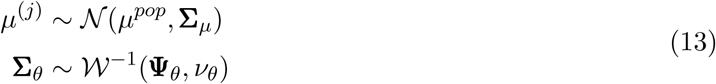

where *µ*^*pop*^ ∈ ℝ^*p*^, *ν*_*θ*_ ∈ ℝ is such that *ν*_*θ*_ > *p*−1, and **Σ**_*µ*_, **Ψ**_*θ*_ are constant, *p*×*p* positive-definite matrices. These choices of hyperpriors are motivated by simplicity as well as computational convenience. Finally, we specify a hyperprior on (*µ*^*pop*^, Σ_*µ*_) of

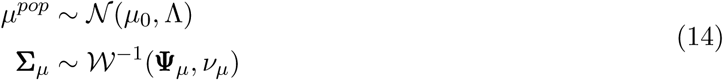

where *µ*_0_ ∈ ℝ^*p*^ and Λ is a constant, *p* × *p* positive-definite matrix.

We set *ν*_*θ*_ = *p, ν*_*µ*_ = *p*, Λ = *I*_*p*_ and use an empirical approach to set the hyperparameters (*µ*_0_, **Ψ**_*µ*_, **Ψ**_*θ*_). We do this by first fitting a single Bayesian ERGM to each network, using the `Bergm` package [8], with the same summary statistics as in the full multilevel model. For each network ***y***^(*i*)^ (ignoring any group membership), we produced *K* = 15000 samples 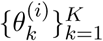 from their respective posterior distributions *π*(*θ*^(*i*)^|***y***^(*i*)^). In particular, we obtained estimates 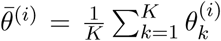 of the posterior mean from each network. Writing 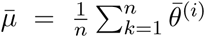, we then set 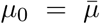 and 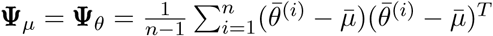.

### A.2 Supplementary results

**Figure A.1:**
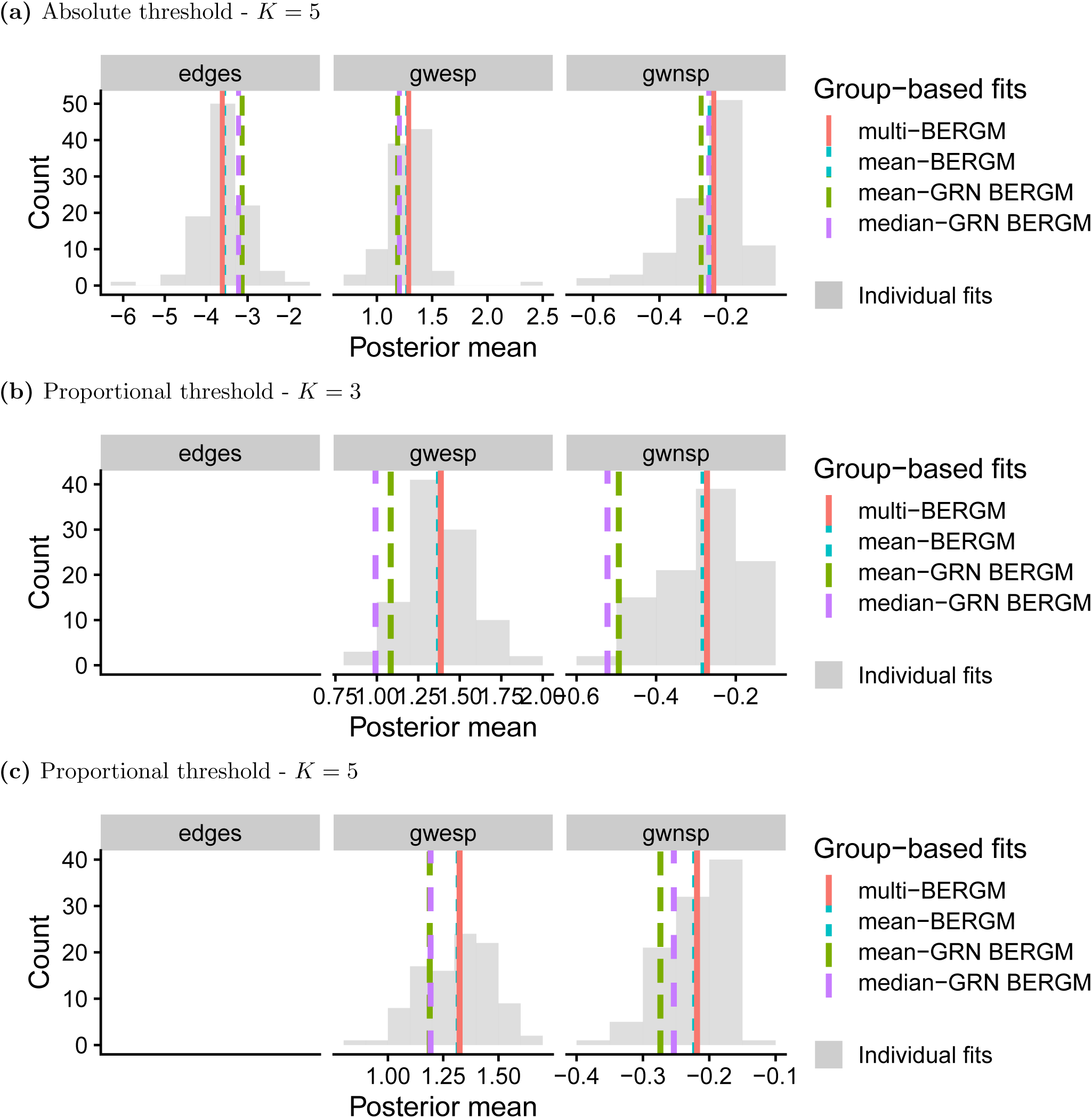
Posterior mean estimates of models fit to networks constructed under a) absolute thresholding with group-wide average node degree *K* = 5, b) proportional thresholding with average node degree *K* = 3 and c) proportional thresholding with average node degree *K* = 5. The histogram (grey bars) corresponds to the posterior mean values of the model parameters fitted to each of the *n* = 100 individual networks. The vertical lines correspond to the posterior mean values of group-level mean parameter under multi-BERGM (red line), mean-BERGM (blue dotted line), mean-GRN BERGM (green dashed line) and median-GRN BERGM (purple dashed line). Under absolute thresholding with average node degree *K* = 5, the posterior mean estimates were broadly similar for each approach. Under proportional thresholding, however, while the posterior mean under multi-BERGM lay close to the mean-BERGM estimate, i.e. the centre of the single-BERGM mean values the posterior means for the mean-GRN BERGM and median-GRN BERGM lay more in the tails of the single-BERGM mean values.

**Figure A.2:**
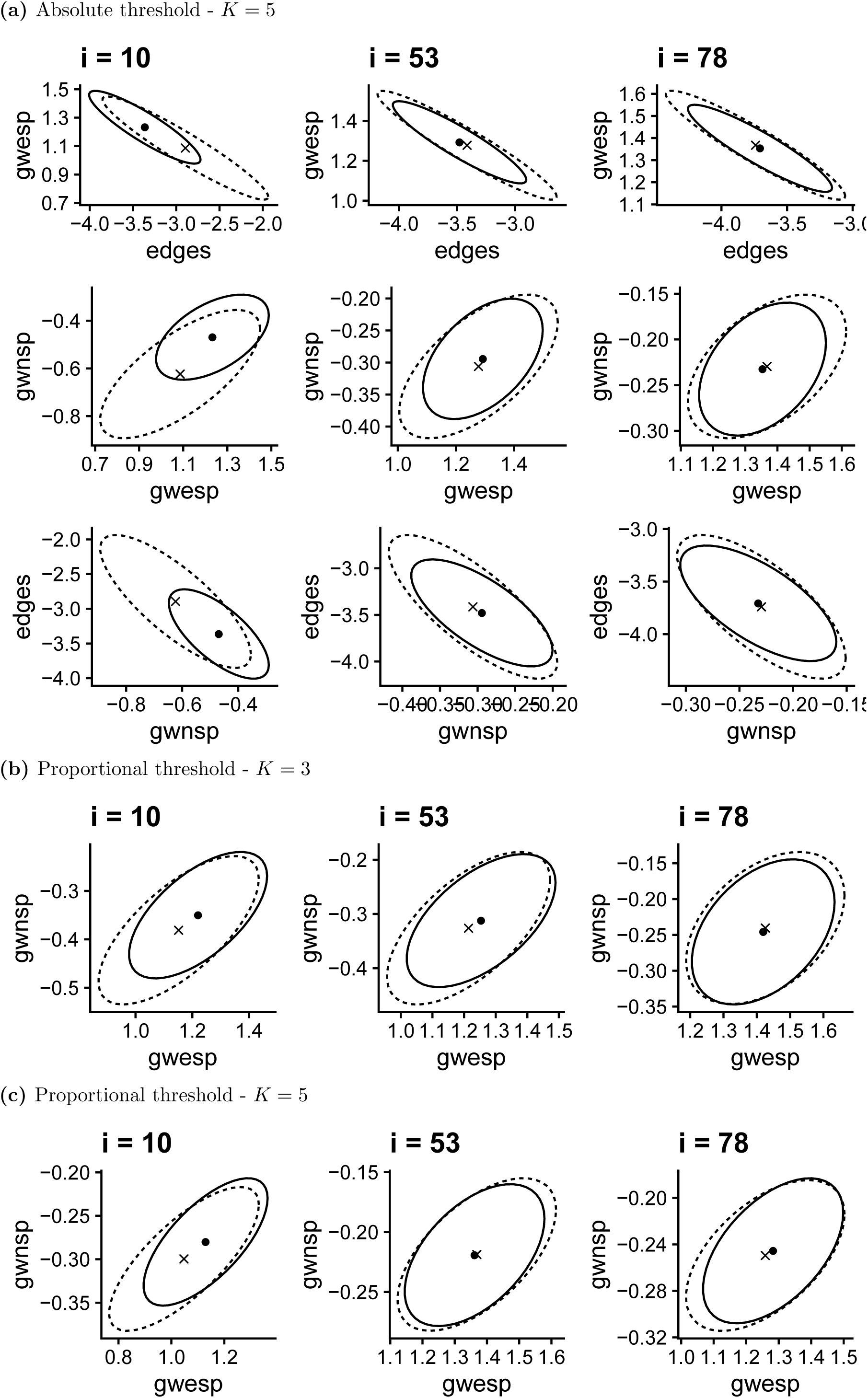
Posterior means and 95% credible regions of the individual-level parameters for three subjects and for each pair of model components under the multi-BERGM (dot and solid line) and under the single-BERGM (cross and dashed line) approaches.

**Figure A.3:**
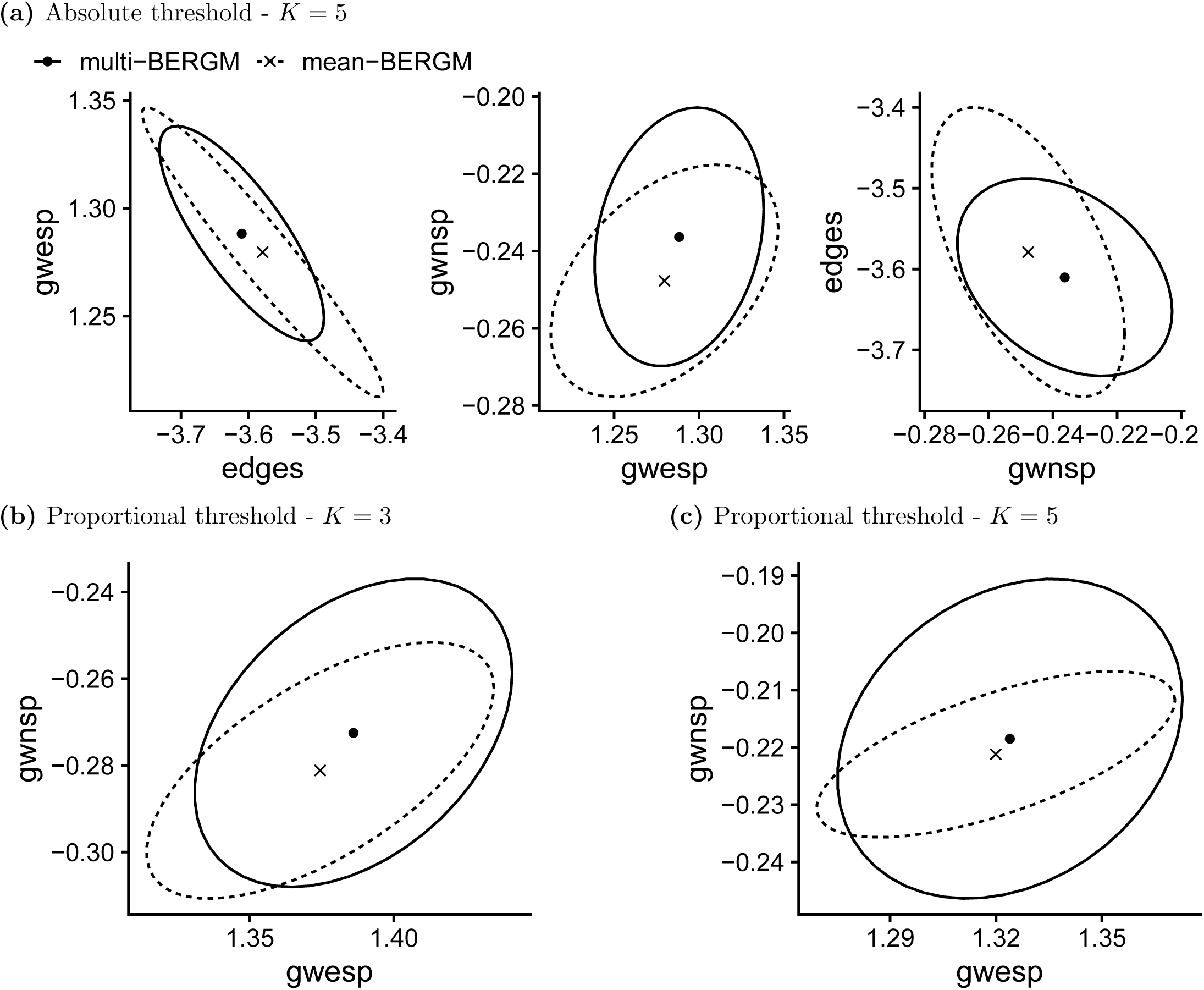
Posterior means and 95% credible regions of the group-level mean parameters for each pair of model components under the multi-BERGM (dot and solid line) and under the mean-BERGM (cross and dashed line) approaches.

**Figure A.4:**
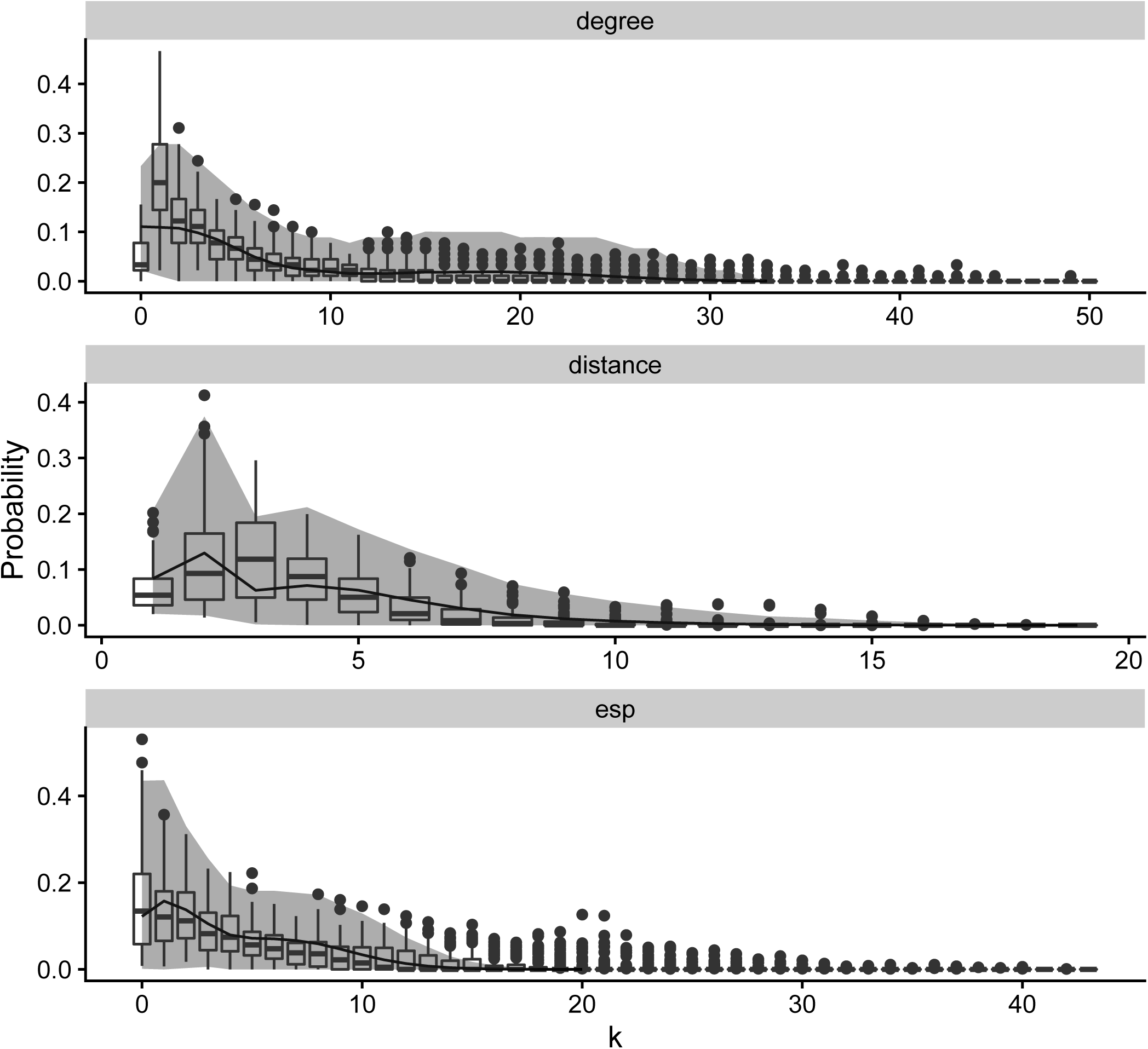
Absolute threshold with group-wide average node degree *K* = 5. Goodness-of-fit assessment of the multi-BERGM for a single group. *S* = 200 networks were generated from the ERGM with model parameters sampled from the respective group-level distributions. The simulated networks are compared to the observed data (shown in the box plots) across three network metrics: degree distribution, geodesic distance distribution, edge-wise shared partner distribution. The lines correspond to the respective means across the simulated networks, while the ribbons correspond to 95% credible intervals.

**Figure A.5:**
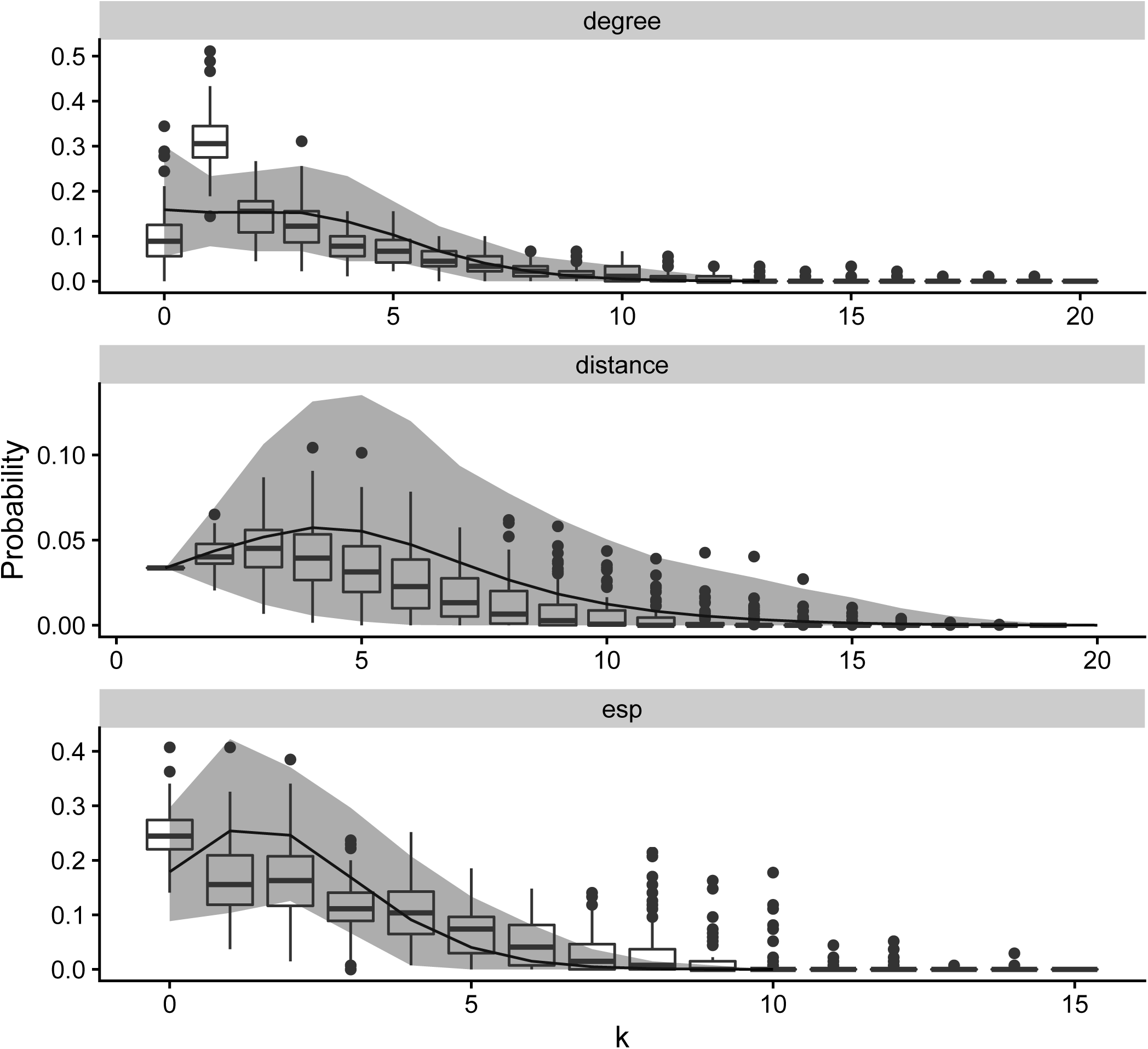
Proportional threshold with average node degree *K* = 3. Goodness-of-fit assessment of the multi-BERGM for a single group. *S* = 200 networks were generated from the ERGM with model parameters sampled from the respective group-level distributions. The simulated networks are compared to the observed data (shown in the box plots) across three network metrics: degree distribution, geodesic distance distribution, edge-wise shared partner distribution. The lines correspond to the respective means across the simulated networks, while the ribbons correspond to 95% credible intervals.

**Figure A.6:**
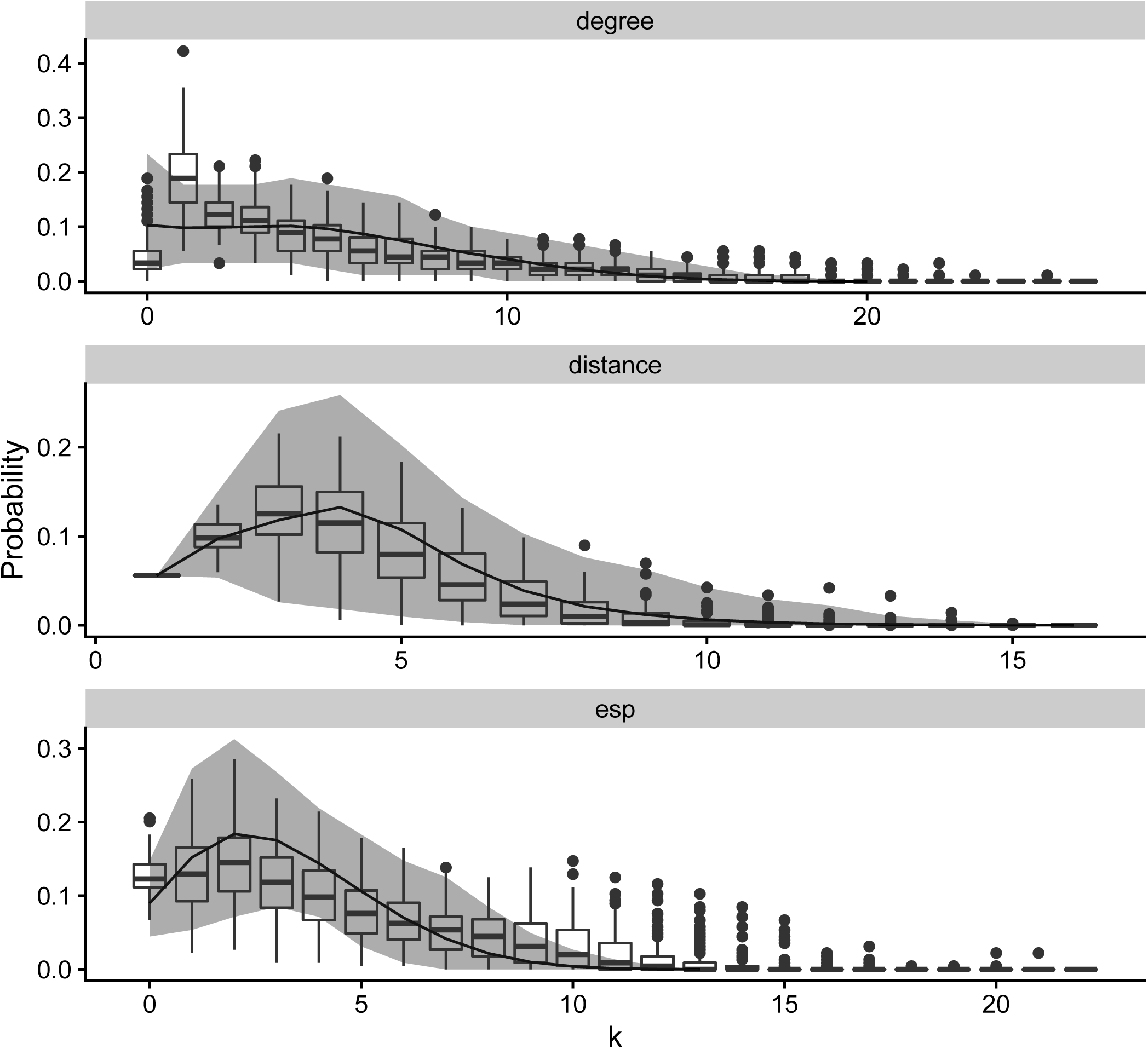
Proportional threshold with average node degree *K* = 5. Goodness-of-fit assessment of the multi-BERGM for a single group. *S* = 200 networks were generated from the ERGM with model parameters sampled from the respective group-level distributions. The simulated networks are compared to the observed data (shown in the box plots) across three network metrics: degree distribution, geodesic distance distribution, edge-wise shared partner distribution. The lines correspond to the respective means across the simulated networks, while the ribbons correspond to 95% credible intervals.

**Figure A.7:**
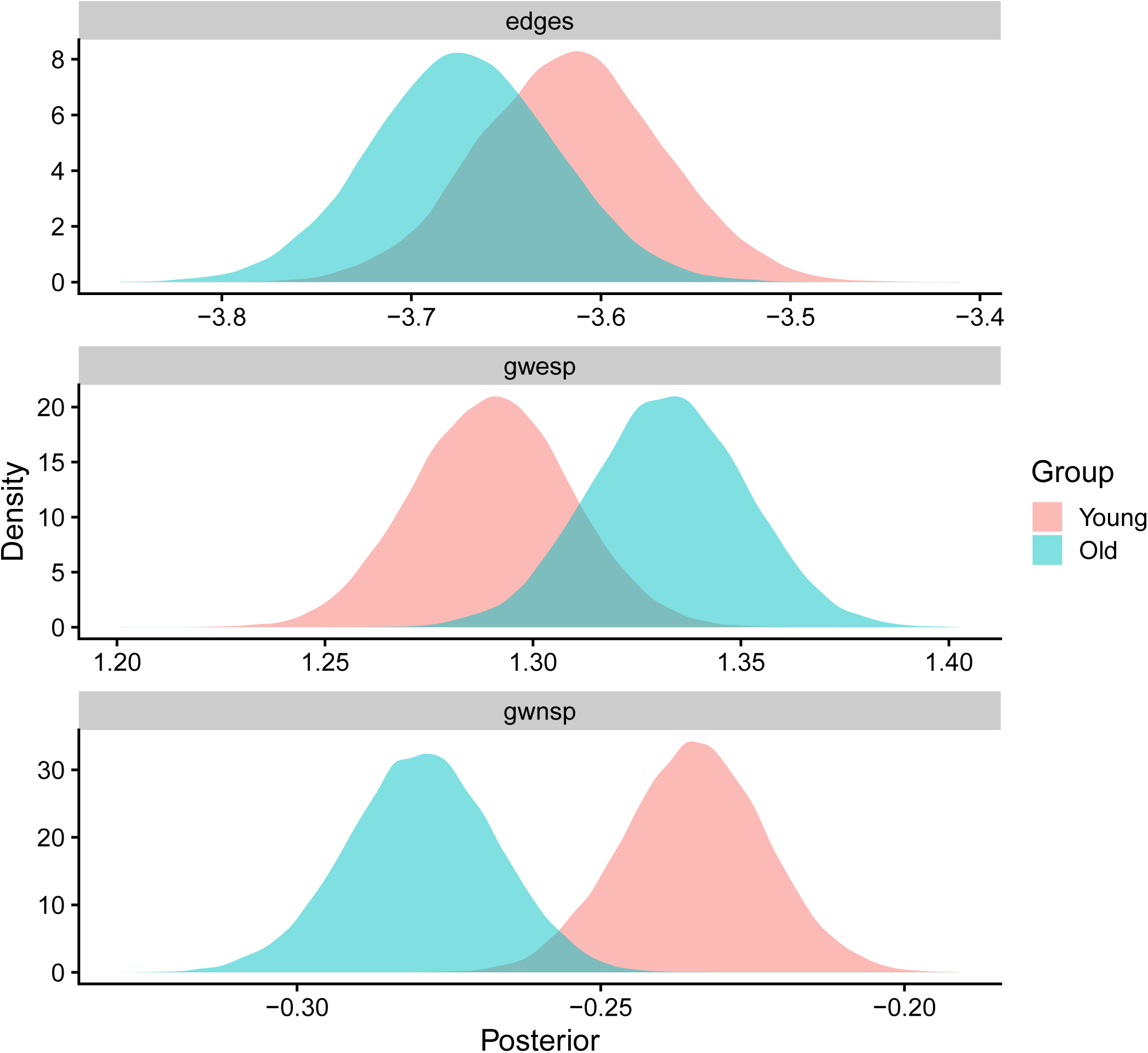
Posterior density estimates under multi-BERGM for the group-level mean parameters for networks constructed via absolute thresholding with average node degree *K* = 5. The edges and GWNSP parameters were moderately larger for the young group while the GWESP parameter was larger for the old group.

**Figure A.8:**
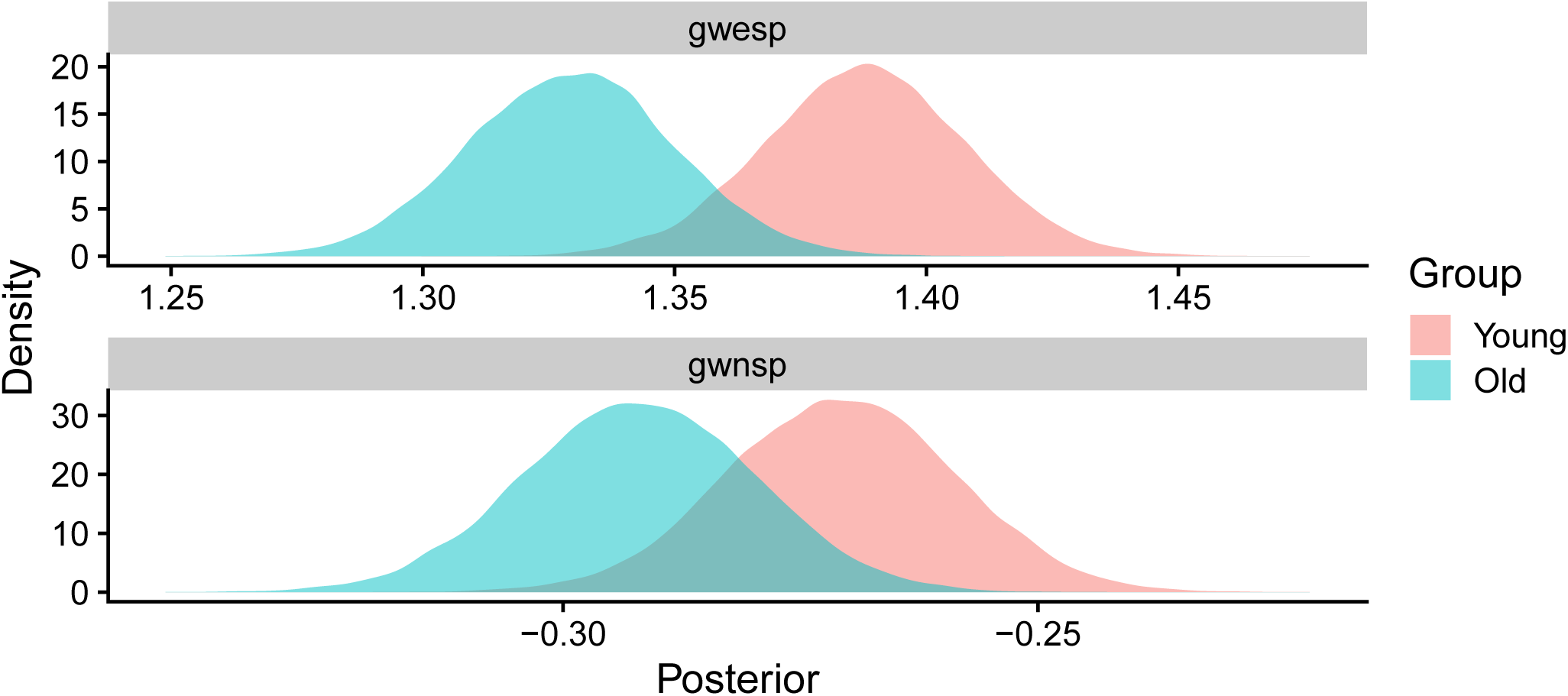
Posterior density estimates under multi-BERGM for the group-level mean parameters for networks constructed via proportional thresholding with average node degree *K* = 3. Both the GWESP and GWNSP parameters were moderately larger for the young group.

**Figure A.9:**
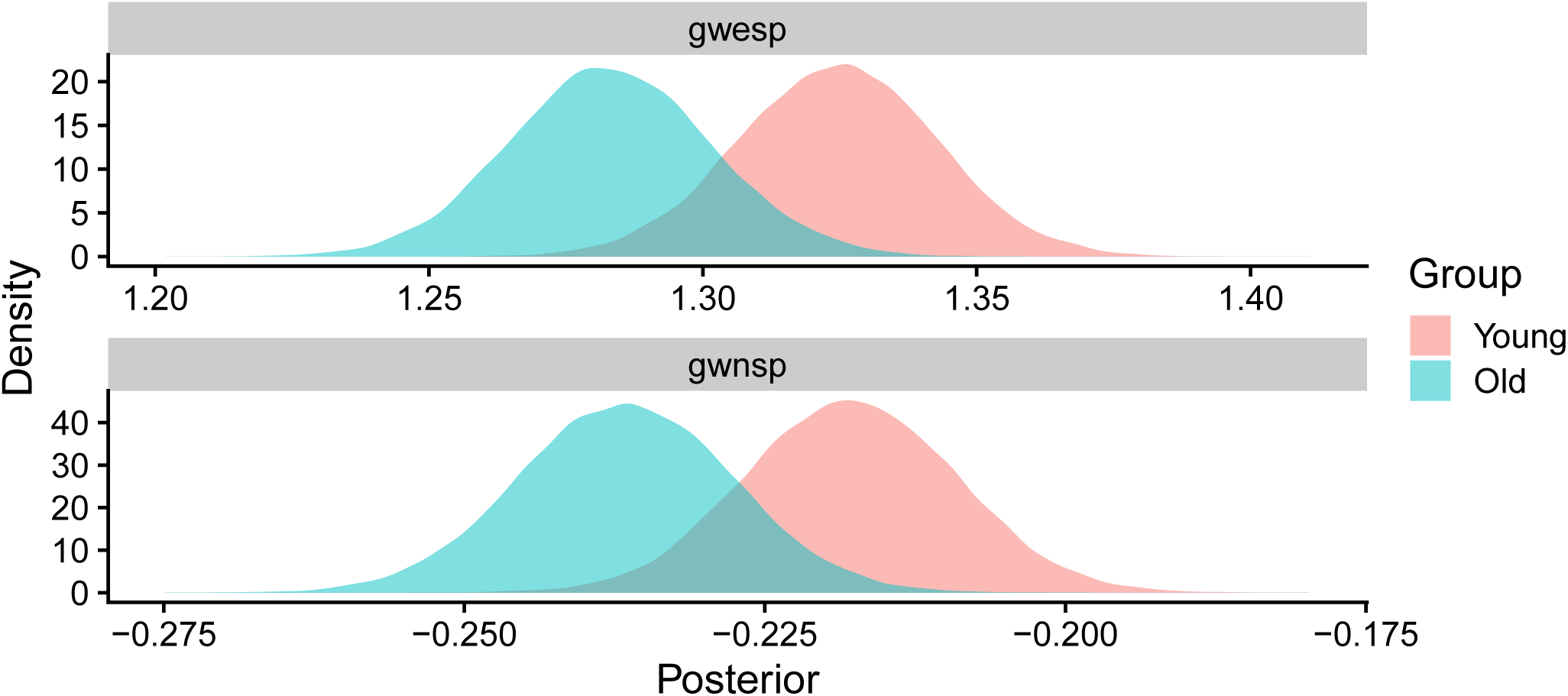
Posterior density estimates under multi-BERGM for the group-level mean parameters for networks constructed via proportional thresholding with average node degree *K* = 5. Both the GWESP and GWNSP parameters were moderately larger for the young group.

**Figure A.10:**
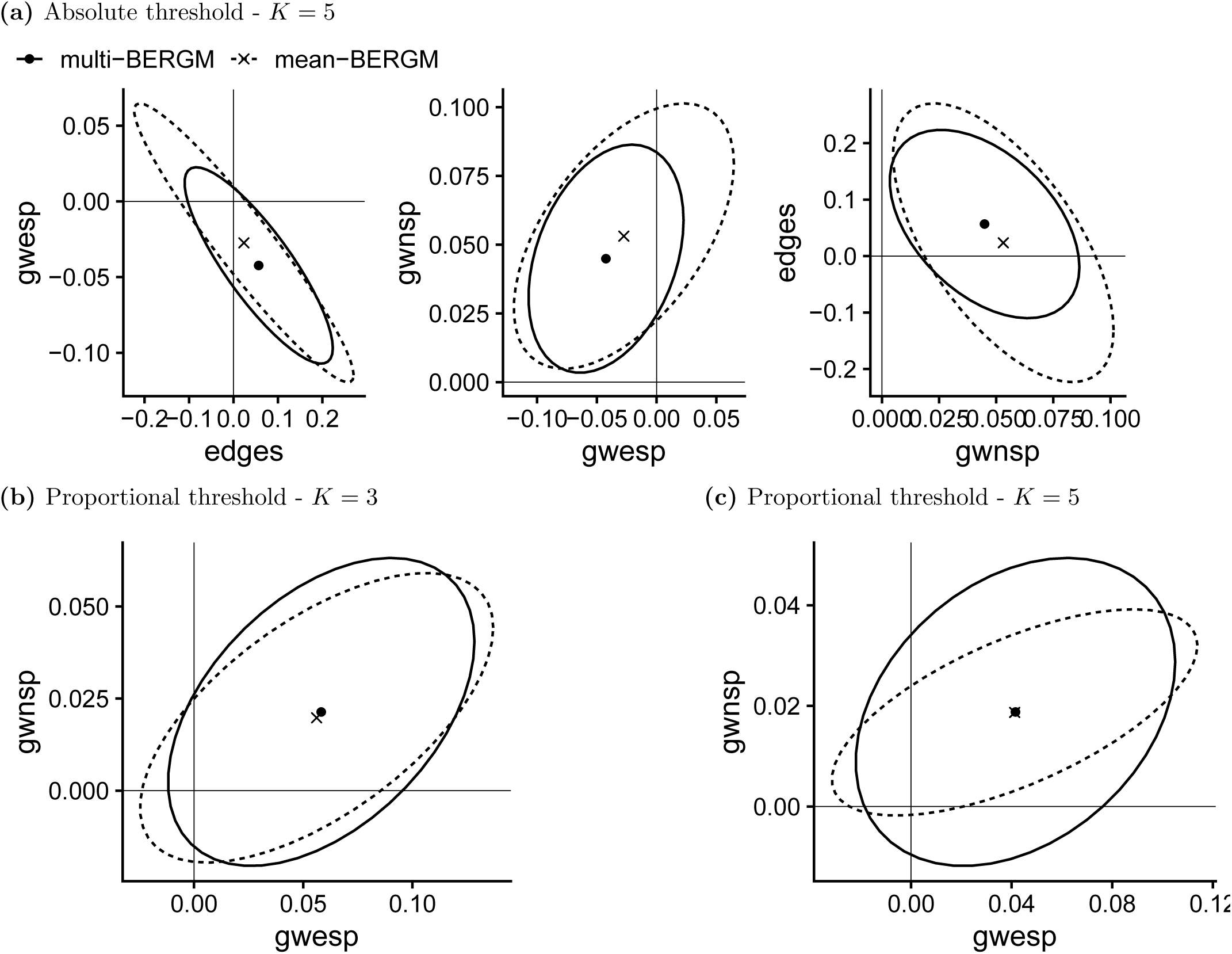
Posterior means and 95% credible regions of the difference in group-level mean parameters for each pair of model components under the multi-BERGM (dot and solid line) and under the mean-BERGM (cross and dashed line) approaches.

**Figure A.11:**
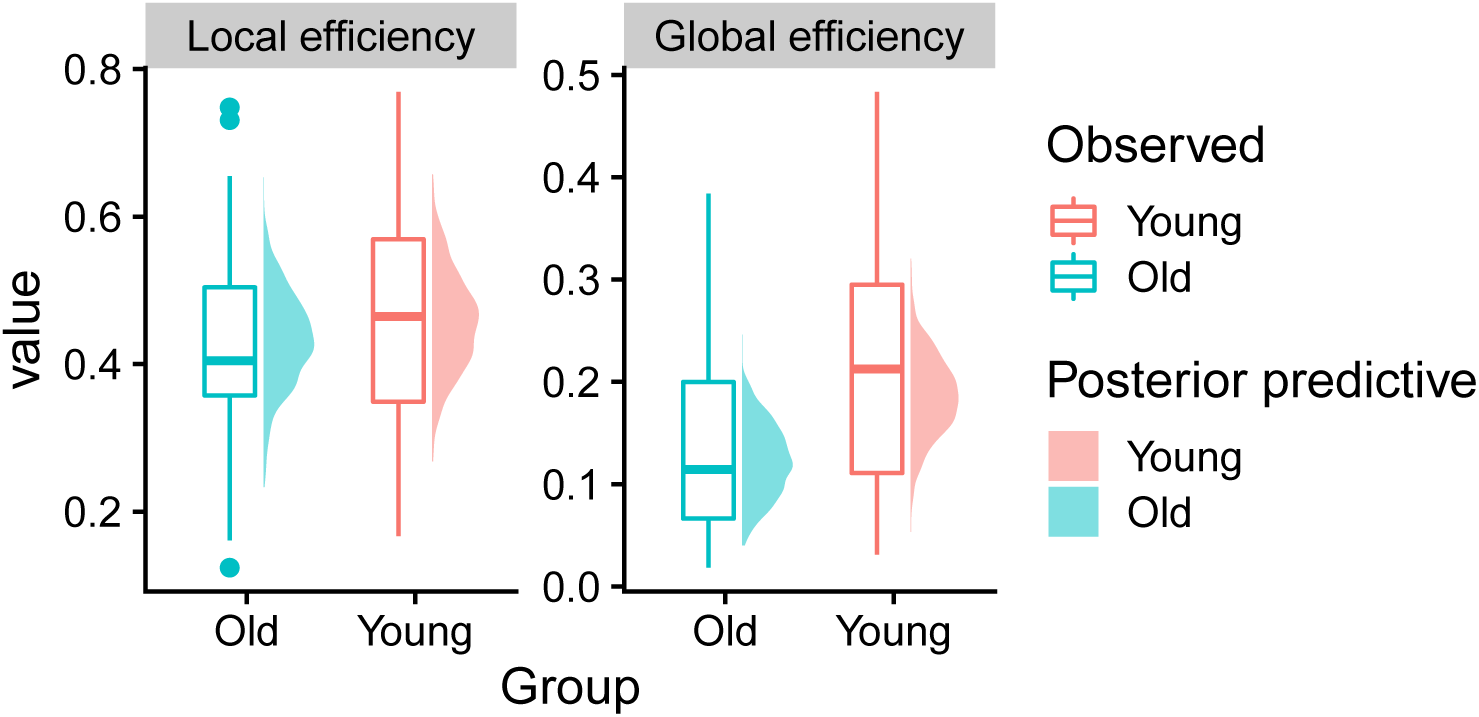
Local and global efficiency in the observed networks constructed via absolute thresholding with average node degree *K* = 5 (bar plots) compared to *S* = 1000 networks simulated from the posterior predictive distribution (density plots) for the young group (red) and the old group (blue).

**Figure A.12:**
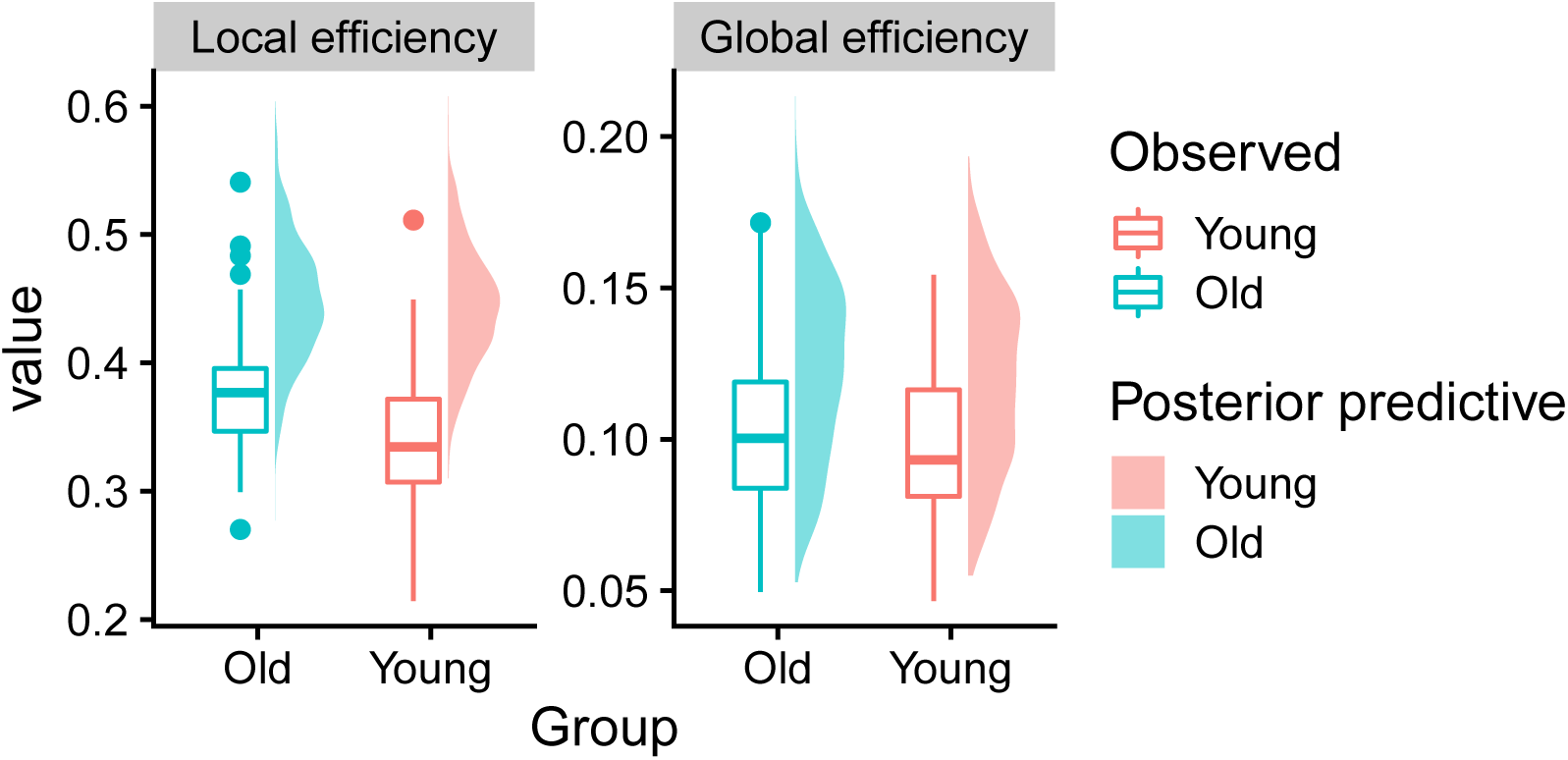
Local and global efficiency in the observed networks constructed via proportional thresholding with average node degree *K* = 3 (bar plots) compared to *S* = 1000 networks simulated from the posterior predictive distribution (density plots) for the young group (red) and the old group (blue).

**Figure A.13:**
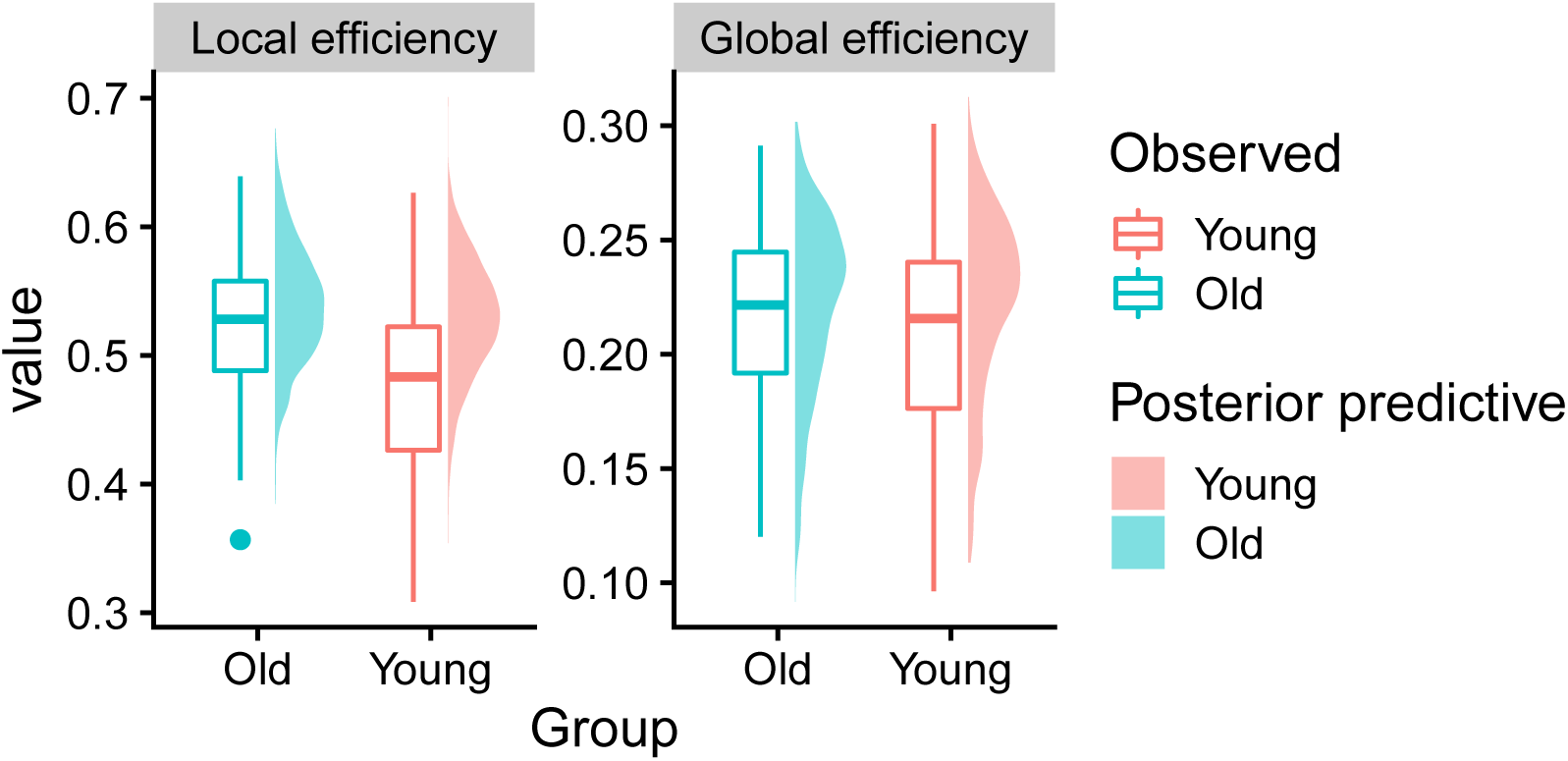
Local and global efficiency in the observed networks constructed via proportional thresholding with average node degree *K* = 5 (bar plots) compared to *S* = 1000 networks simulated from the posterior predictive distribution (density plots) for the young group (red) and the old group (blue).

